# Profiling the neuroimmune cascade in 3xTg mice exposed to successive mild traumatic brain injuries

**DOI:** 10.1101/2023.06.13.544838

**Authors:** Alyssa F. Pybus, Sara Bitarafan, Rowan O. Brothers, Alivia Rohrer, Arushi Khaitan, Felix Rivera Moctezuma, Kareena Udeshi, Brae Davies, Sydney Triplett, Eric Dammer, Srikant Rangaraju, Erin M. Buckley, Levi B. Wood

## Abstract

Repetitive mild traumatic brain injuries (rmTBI) sustained within a window of vulnerability can result in long term cognitive deficits, depression, and eventual neurodegeneration associated with tau pathology, amyloid beta (Aβ) plaques, gliosis, and neuronal and functional loss. However, we have limited understanding of how successive injuries acutely affect the brain to result in these devastating long-term consequences. In the current study, we addressed the question of how repeated injuries affect the brain in the acute phase of injury (<24hr) by exposing the 3xTg-AD mouse model of tau and Aβ pathology to successive (1x, 3x, 5x) once-daily weight drop closed-head injuries and quantifying immune markers, pathological markers, and transcriptional profiles at 30min, 4hr, and 24hr after each injury. We used young adult mice (2-4 months old) to model the effects of rmTBI relevant to young adult athletes, and in the absence of significant tau and Aβ pathology. Importantly, we identified pronounced sexual dimorphism, with females eliciting more differentially expressed proteins after injury compared to males. Specifically, females showed: 1) a single injury caused a decrease in neuron-enriched genes inversely correlated with inflammatory protein expression as well as an increase in AD-related genes within 24hr, 2) each injury significantly increased expression of a group of cortical cytokines (IL-1α, IL-1β, IL-2, IL-9, IL-13, IL-17, KC) and MAPK phospho-proteins (phospho-Atf2, phospho-Mek1), several of which were co-labeled with neurons and correlated with phospho-tau, and 3) repetitive injury caused increased expression of genes associated with astrocyte reactivity and immune function. Collectively our data suggest that neurons respond to a single injury within 24h, while other cell types including astrocytes transition to inflammatory phenotypes within days of repetitive injury.

## INTRODUCTION

Traumatic brain injury (TBI) results in approximately 2.5 million emergency department visits every year in the Unites States alone. Approximately 75% of TBI cases are classified as mild (mTBI), and these mild cases carry an annual cost burden of $17 billion (1,2). An estimated 10-40% of mTBIs are associated with persistent cognitive impairment lasting longer than one month and in some cases up to a year (3–6). Moreover, repeated mTBIs (rmTBIs) sustained within a window of vulnerability can intensify pathological and functional consequences (7,8). These repeated injuries, which are often observed among athletes in high contact sports such as American football and boxing, can result in the development of Alzheimer’s disease-like pathology, including neurofibrillary tangles and amyloid beta (Aβ) plaques (9). Current treatment paradigms for (r)mTBI are severely lacking and focus on the alleviation of symptoms, rather than targeting the underlying mechanisms of injury. There is thus an urgent need to illuminate the post-injury neuro-molecular sequelae for development of targeted therapies.

Mounting evidence suggests the involvement of brain immune signaling as a key driver of long-term outcome following mTBI. Brain immune signaling is implicated in several neurodegenerative diseases associated with mTBI, including Alzheimer’s disease (AD) (10–12) and Parkinson’s disease (PD) (13–15). Moreover, the roles of immune signaling in driving pathology following severe TBI have been relatively well-studied (16–20), and preliminary evidence shows similar patterns in (r)mTBI (16,21–25). Indeed, prior research from our own group has identified correlations between lower cerebral blood flow (a potential biomarker of worse cognitive outcome) and increased mitogen-activated protein kinase (MAPK) signaling, cytokine expression, and microglial activation after repetitive closed-head injuries (CHI) in wild type mice, supporting an essential relationship between brain immune signaling and outcome after mTBI. However, a comprehensive study relating acute changes in immune signaling and glial activation to neuronal changes and pathological markers after single and repetitive mTBIs is currently lacking and would provide a much-needed perspective to guide the search for new therapies.

The acute pathophysiology of mTBI has previously been defined by the “neurometabolic cascade of concussion,” in which neurons undergo axonal injury and dysfunction, altered neurotransmission, ionic flux, and indiscriminate release of glutamate, while many brain cells undergo a hyperglycolysis energy crisis to restore homeostasis (26,27). When the neurometabolic cascade was defined in 2014, relatively little was known about mTBI-induced inflammatory changes, but research over the last decade has more fully defined changes in immune signaling, microglial activation, and astrocyte reactivity, as well as their effects on secondary injury after TBI (16,28–31). For example, recent studies suggest that astrocytes rapidly respond to mechanical stress after injury, potentially via opening of mechanosensitive ion channels, leading to activation of MAPK signaling, ATP release, and reactive astrogliosis (28,32–38). Here we propose to similarly define the acute “neuroimmune cascade” of mTBI. These clear changes in microglial and astrocyte responses post-mTBI suggest a broad and complex neuroimmune response to mTBI. Thus, a comprehensive understanding of neuroimmune signaling and its hypothesized relationship to neuronal, glial, and pathogenic changes after injury is critical to identifying drivers of secondary injury as well as determining therapeutic targets to mitigate adverse cognitive and pathological outcomes.

In the current study, we sought to define the neuroimmune cascade of rmTBI by profiling protein and transcriptomic changes associated with immune signaling, pathology, and the neuronal, microglial, and astrocytic compartments. We specifically hypothesized that mTBI would drive brain immune signaling in a manner correlated with markers of neurodegeneration, such as tau, Aβ, astrocyte reactivity, and microglial activation. Because wild-type mice show inconsistent evidence of pathological changes after TBI (39–41), we used the triple transgenic model of Alzheimer’s-like pathology (3xTg-AD), which contains human mutant forms of genes in the Aβ processing pathway (*APP*, *PSEN1*) and tau (*MAPT*). This model is widely used to study AD and is increasingly utilized in brain injury research (42–49). Here, we sought to concomitantly define the effects of successively increasing mild traumatic brain injuries on the following molecular outcomes in the frontal cortex and hippocampus: 1) the kinetics of brain immune signaling and cytokine expression, 2) tau phosphorylation and Aβ burden, 3) markers of astrocyte reactivity and microglial activation, and 4) transcriptional profiling of the somatomotor cortex (females only). Our data analyses revealed pronounced effects of repetitive injury on all molecular outcomes. Further, the inclusion of both male and female 3xTg-AD mice in our study revealed strong effects of sexual dimorphism in this model across all outcomes, with males showing higher basal levels of immune signaling but females showing more pronounced changes in response to injury.

## MATERIALS AND METHODS

### Study Protocol

3xTg-AD on C57BL/6J mice (Jackson Laboratory, strain 033930) were aged 2-4 months (98 females, 86 males). All protocols were approved by the U.S. Army Medical Research and Development Command Animal Care and Use Review Office (ACURO) and the Emory University Institutional Animal Care and Use Committee (IACUC) in accordance with National Institutes of Health guidelines. Mice were housed in a pathogen free facility with a twelve-hour light/dark cycle. Food and water were provided *ad libitum*. Mice were randomly assigned to one of four injury groups: sham-injured controls, one closed head injury (1xCHI), three closed-head injuries spaced once daily (3xCHI), and five closed-head injuries spaced once daily (5xCHI). Animals were sacrificed, and the brain was harvested for molecular and pathological assessment at the following time points for each group: pre-injury, 30 minutes, 4 hours, and 24 hours post-injury. Sample sizes (n=6-8 per injury time point group in protein data, n=2-4 in RNAseq data) are provided in **Table S1**.

### Closed Head Injury Model

This work was conducted using a previously characterized weight drop closed-head injury (CHI) model (50,51). Animals were anesthetized using 4.5% isoflurane (1 L/min, 70/30 nitric oxide/oxygen) prior to injury. The injury was delivered by dropping a 54 g bolt down a 0.96 m guide tube (49035K85, McMaster-Carr, Elmhurst IL) onto the dorsal aspect of the head (skull intact), targeting between the approximate location of the coronal and lambdoid structures. The anesthetized animal was positioned under the guide tube on a Kimwipes® task wipe (Kimberly-Clark, Irving, TX). The animal was grasped by the base of the tail so that on impact, the mouse penetrated the task wipe and underwent rapid, unrestricted rotation of the head in the anterior-posterior plane. Control, sham-injured mice were age- and sex-matched and received the same exposure to anesthesia but were not subject to closed-head injury.

### Quantification of Immune and Pathological Proteins

To comprehensively profile the immune signaling kinetics after repeated CHI in 3xTg-AD male and female mice, we quantified four classes of proteins: glial phenotypic markers (astrocyte reactivity markers GFAP and S100B; microglial activation marker CD68 and homeostatic marker TMEM119), molecular markers of pathology (Aβ40, Aβ42, total tau, and phospho-tau T181), nine MAPK phospho-proteins, and 30 cytokines (in total 47 measured proteins, plus ratios Aβ42/40 and pTau/tTau). We collected frontal cortex and hippocampal samples from the left hemisphere of the three injury groups (1xCHI, 3xCHI, and 5xCHI) across four time points each (pre-injury and 30min, 4h, 24h post-injury), where the pre-injury 1xCHI group represents sham-injured controls. For analysis purposes, pre-3xCHI and pre-5xCHI are sometimes referred to as the 24h post-2xCHI and 24h post-4xCHI groups, respectively. In total, data was collected from 98 female mice and 86 male mice. Two female and three male mice were excluded from our analysis after Mahalanobis outlier detection using all protein data from the same sex and region (α<0.001), leaving n=96 females and n=83 males.

Cortical and hippocampal brain tissue sections were lysed using the Bio-Plex cell lysis kit (Bio-Rad Laboratories #171304011) and protein concentrations were determined using a Pierce BCA Protein Assay (Thermo Fisher #23225). Multiplexed cytokine quantification was conducted using the Milliplex® MAP Mouse Cytokine/Chemokine 32-Plex kit (Eotaxin, G-CSF, GM-CSF, IFN-γ, IL-1α, IL-1β, IL-2, IL-3, IL-4, IL-5, IL-6, IL-7, IL-9, IL-10, IL-12p40, IL-12p70, IL-13, IL-15, IL-17, IP-10, KC, LIF, LIX, MCP-1, M-CSF, MIG, MIP-1α*, MIP-1β, MIP-2, RANTES, TNF-α, and VEGF*) (Millipore Sigma MCYTMAG-70K-PX32). Cytokines marked with an asterisk did not fall within a linear range and were not included in our analysis. Multiplexed phospho-protein quantification was conducted for the MAPK signaling pathway using the Bio-Plex Pro™ Cell Signaling MAPK Panel 9-plex kit (phospho-Atf2 T71, phospho-Erk1/2 T202/Y204 T185/Y187, phospho-HSP27 S78, phospho-JNK T183/Y185, phospho-Mek1 S217/S221, phospho-p38 MAPK T180/Y182, phospho-p53 S15, phospho-p90 RSK S380, phospho-Stat3 S727) (Bio-Rad Laboratories LQ00000S6KL81S). Prior to analysis, lysates were thawed on ice and centrifuged at 4°C for 10 min at 15,500g. Protein concentrations were normalized with Milliplex® MAP Assay Buffer (EMD Millipore, Billerica, MA) to 6 μg protein per 12.5 μL for cytokine analysis and 1 μg protein per 12.5 μL for the MAPK pathway analysis. These protein concentrations were selected because they fell within the linear range of bead fluorescent intensity vs. protein concentration for detectable analytes. Quantification of pathological markers was conducted for total tau, phospho-tau (T181), Aβ 1-40, and Aβ 1-42 using the Milliplex® MAP Human Amyloid Beta and Tau Multiplex Assay kit (Millipore Sigma HNABTMAG-68K). All kits were read on a MAGPIX® system (Luminex, Austin, TX).

We quantified astrocyte reactivity markers glial fibrillary acidic protein (GFAP) and S100 calcium binding protein B (S100B) as well as macrophage and microglia activation marker cluster of differentiation 68 (CD68) and microglial homeostatic marker transmembrane protein 119 (TMEM119) via enzyme-linked immunosorbent assay (ELISA). Protein concentrations were normalized with respective assay diluents from each kit to 2.5 μg/ml in 50 μL for the GFAP ELISA kit (Abcam ab233621), 50 μg/ml in 50 μL for the S100B ELISA kit (Abcam ab234573), and 0.1 μg/μL in 50ul for the CD68 and TMEM119 ELISA kits (Lifespan Biosciences LS-F11095 and LS-F52734). These protein concentrations were selected because they fell within the linear range of absorbance vs. protein concentration for detectable analytes.

### Data Analysis and Visualization

Data was analyzed and figures were generated in RStudio (Boston, MA) using the R programming language. Data processing was conducted using the `tidyverse` collection of packages (52). Heatmaps were generated using the R package `heatmap3` (53), bar graphs and regression plots were created using the packages `ggplot2` (54) and `ggpubr` (55), and gene set variation analysis was conducted using the `gsva` package (56) available on Bioconductor. For each molecular assay, we collected background measurements using assay buffer in the absence of biological samples. We subtracted average background measurements from each sample measurement and set negative values to zero. Clustering was conducted using the `hclust` function of the `stats` package in R using Euclidean distance in the unweighted pair group method with arithmetic mean. Outlier detection was conducted in R by calculating Mahalanobis distance of each point from the data’s centroid within each grouping of sex and region (female frontal cortex, female hippocampus, male frontal cortex, and male hippocampus) using the `ClassDiscovery` (57) package. Two female and three male hippocampus samples were outside the cutoff threshold of α=0.001 and all data from those animals was discarded in both the frontal cortex and hippocampus sets.

### Adjustment for Batch Effect and Pathology

Male and female data were collected simultaneously for S100B, CD68, and TMEM119 by ELISA, but separated into different batches for all other protein measurements. Eight female sample replicates were included in each male batch assay and then used to transform the male data into the female space via a linear model for each protein. Computations were conducted using the lm() function in R.

To adjust data for the effects of pathology, linear models were calculated using neuroimmune proteins as dependent variables with the following combinations of independent variables: total tau and phospho-tau T181; Aβ40 and Aβ42; or total tau, phospho-tau T181, Aβ40, and Aβ42. Residuals of each linear model were defined as the pathology-adjusted (tau-adjusted, Aβ-adjusted, or total pathology-adjusted) neuroimmune protein data. Computations were conducted using the lm() function in R.

### Immunohistochemistry

Immunohistochemistry was performed to investigate GFAP, NeuN, IL-1α, IL-1β, IL-13, KC, phospho-Atf2 (T71), phospho-Mek1/2 (S217/S221), and phospho-tau (S262/T263). Right brain hemispheres from each animal were fixed in 4% paraformaldehyde then processed and embedded in paraffin. Tissue slices were cut into 10 µm thick sagittal sections using a rotary microtome (Thermo Fisher) and affixed onto glass microscopy slides (Electron Microscopy Sciences, Hatfield, PA). Tissue slices were deparafinized in xylene and rehydrated by washing with 100% ethanol, 95% ethanol, and deionized water. Antigen retrieval was performed in a microwave by boiling slides in 10mM sodium citrate buffer at pH 6.0. Slides were rinsed in tris-buffered saline with 0.01% tween (TBST). A hydrophobic ring was drawn around each individual tissue slice using an immunohistochemistry PAP pen (Enzo Life Sciences, Farmingdale, NY), after which samples were blocked for 2 hr in blocking buffer, 5% w/v Bovine Serum Albumin (BSA) or 50% goat serum (Sigma-Aldrich) in TBST. Samples were incubated at 4°C overnight with primary antibodies diluted in blocking buffer: GFAP (1:100, Diagnostic Biosystems Mob064), NeuN (1:200, Abcam ab104224), IL-1α (1:50, Thermo Fisher PA5-89037), IL-1β (1:500, Abcam ab283818), IL-13 (1:100, Abcam ab106732), KC (1:50, Thermo Fisher PA5-86508), phospho-Atf2 T71 (1:50, Thermo Fisher PA5-97332), phospho-Mek1/2 S217/S221 (1:50, Themo Fisher PA5-1057), or phospho-tau S262/T263 (1:100, Abcam ab92627), as appropriate. Slides were rinsed again in TBST and incubated with either Alexa Fluor 555 and/or Alexa Fluor 488 secondary antibodies (Thermo Fisher) diluted 1:200 in blocking buffer. Slides were counterstained with 1 µg/mL DAPI, rinsed in water, and mounted with VECTASHIELD Antifade Mounting Medium (Vector Laboratories, Burlingame, CA). Samples were imaged using epifluorescent microscopy on a Zeiss Axio Observer Z.1 inverted microscope.

### Bulk Tissue RNA Sequencing and Analysis

We conducted RNA sequencing on a representative subset of 40 female left somato-motor cortex samples selected from the original 96 for bulk tissue RNA sequencing (available on the NCBI Gene Expression Omnibus under accession number GSE226838). One sample was removed post-sequencing after identification as an outlier animal within the protein data. RNA was extracted using the miRNeasy Micro kit (Qiagen #217084). Extracted RNA was sent to Admera Health, LLC (South Plainfield, NJ) for sequencing, alignment, and calculation of the count matrix. The count matrix yielded 32,807 non-zero features. Features with fewer than 10 counts in more than four samples (∼10%) were filtered out of the analysis (17,844 features). The remaining 14,963 features were normalized by ratio of median method then variance-stabilized in R using the `DESeq2` package, available on Bioconductor (58).

Weighted gene co-expression network analysis (WGCNA) was conducted in R using the `WGCNA` package (59) available on the Comprehensive R Archive Network (CRAN). A WGCNA threshold power of 4 was chosen since it was the smallest threshold that resulted in a scale-free R^2^ value greater than 0.80. We constructed our network in a single block using blockwiseModules() with the following parameters: threshold power of 4, signed modules, a minimum module size of 20, a merge cut height of 0.07, a reassignment threshold of 0.05, a “bicor” correlation type, and a “mean” TOM denominator. Significance of module eigengene (ME) expression between sham and injured samples was assessed by permutation test, in which ME values for both populations were randomly re-assigned to either group then permutated mean difference was compared to the true mean difference across 10,000 iterations. Original sample sizes were maintained. Groups were considered significantly different if the true mean difference was greater than or equal to the permutated mean difference in at least 95% of iterations.

Gene ontology was conducted for each of the 14 resulting WGCNA modules using PANTHER overrepresentation test on PANTHER 17.0, available on the Gene Ontology Resource. We used the mm10 GO biological process complete annotation set with Fisher’s exact test and false discovery rate (FDR) corrected p-values. The test was applied using all genes with a network kME, or module connectivity (59), of at least 0.60 and a background of the 14,963 features used to construct the network.

Cell type enrichment was conducted using protocol as published in (60–62). Briefly, a cell type marker list was created for neurons, oligodendrocytes, endothelia, astrocytes, and microglia from cell type specific mouse brain proteome and transcriptome studies Sharma *et al.* (63) and Zhang *et al.* (64). Fisher exact tests with FDR correction were performed to determine enrichment for each cell type marker list within each WGCNA network module. We used R scripts by Eric Dammer and Divya Nandakumar (Emory University School of Medicine) freely available on GitHub at https://github.com/edammer/CellTypeFET.

Multi-marker Analysis of GenoMic Annotation (MAGMA)(65) was conducted to identify modules with significant enrichment for genes related to Alzheimer’s disease (AD) as identified in genome-wide association studies (GWAS) (66–68). To implement MAGMA, we used R scripts by Eric Dammer as used in Seyfried *et al*. (62), freely available on GitHub at https://github.com/edammer/MAGMA.SPA.

Gene set variation analysis (GSVA) was conducted on the normalized and variance-stabilized count matrix in R using the `gsva` package, also available on Bioconductor (56). Gene sets used for GSVA included the C2 curated gene sets collection from the Molecular Signatures Database (69,70) as well as our previously published astrocyte-enriched gene sets (71).

## RESULTS

### Immune, glial, and pathological markers differed between male and female 3xTg-AD mice

Since prior reports showed pronounced differences in pathological burden between male and female 3xTg-AD mice (72,73), we began the current study by evaluating sex differences among all protein measures: cytokines, MAPK phospho-proteins, glial markers, phosphorylated and total tau, and Aβ40/42. Data were adjusted for batch effects as needed using linear models of technical replicates (**Methods**). We first obtained a holistic view of variation within the data using a principal component analysis (PCA), which revealed a pronounced effect of sex within both brain regions (**Fig. 1A**). Female replicate samples from the male batches grouped with female batch samples and separately from male samples, emphasizing that the effects of sex were biological, rather than technical (purple points, **Fig. 1A**).

**Figure 1.**
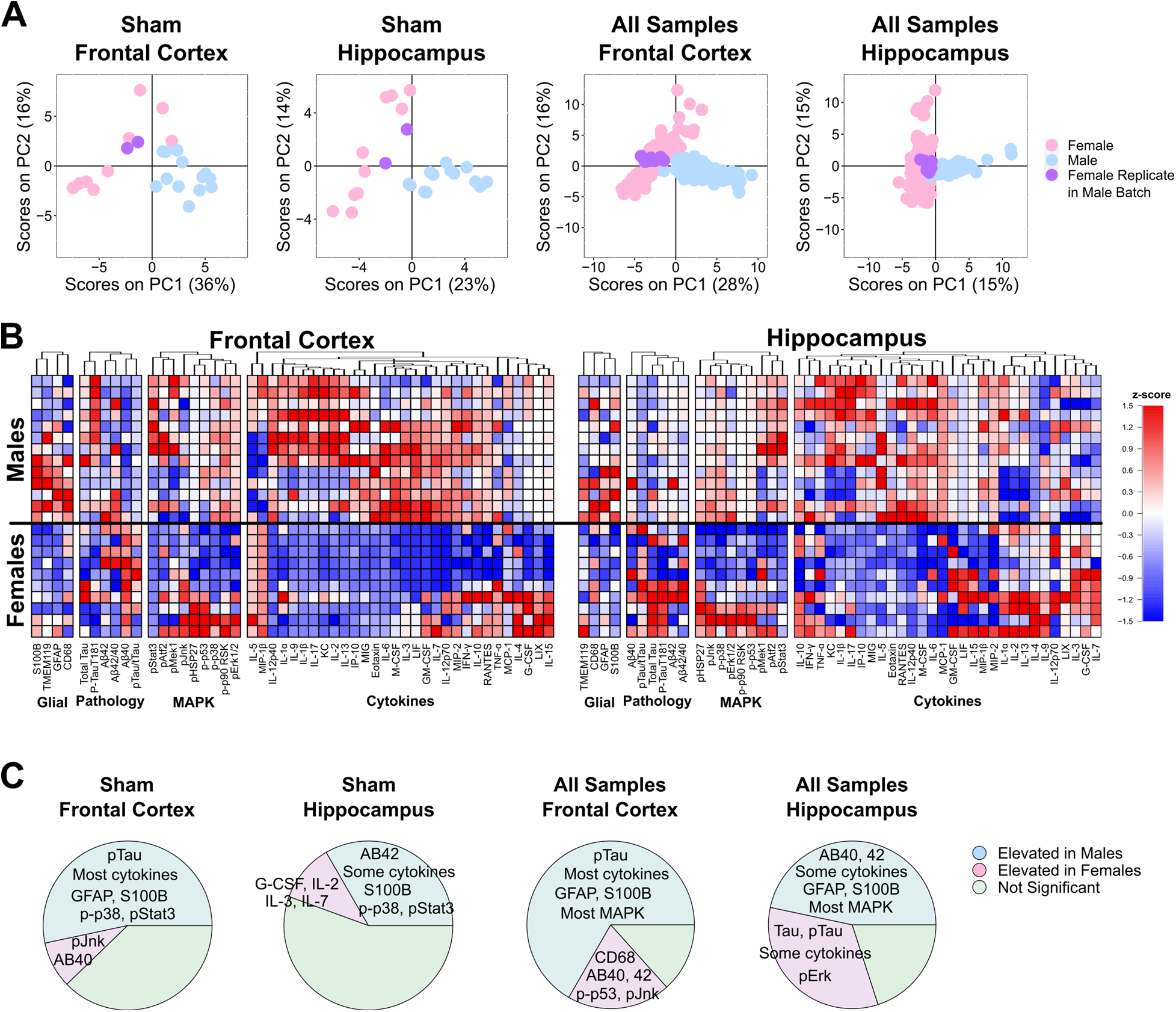
Pronounced sexual dimorphism of immune signaling, glial markers, and molecular markers of pathology in 3xTg-AD mice. **A**) Principal component analysis of combined protein data across sham-injured animals (left) and all injuries and time points (right) reveals strong separation between male and female mice on the first two PCs. Female replicate samples measured in male batch assays are shown in purple. Percent variance described by each PC is annotated in parentheses. **B**) Uninjured control mice show distinct baseline sex differences in expression of glial markers, pathological markers, MAPK signaling, and cytokines. Proteins are categorized by function then clustered by Euclidean distance within each category. Red to blue indicates relatively high to low expression among all sham animals (n=13 males, n=10 females). Of the 46 measured immune and pathological marker proteins in sham-injured animals, 37 from the frontal cortex (80%) and 30 from the hippocampus (65%) had higher average expression in males compared to females. **C**) Pie charts for each region and sample set (just sham on left, all animals combined on right) show high proportions of all analytes measured (4 glial markers, 4 pathological markers, 9 MAPK, 30 cytokines) that are significantly elevated (p<0.05, Wilcoxon rank-sum) in males (blue) or females (pink). See summary of significant differences in **Table S2**.

To define the specific effects of sex versus injury and time point, we conducted multiple linear regression of all quantified proteins and phospho-proteins using sex, injury number, and time point as independent variables, revealing that sex, as compared to all other variables in the model, had the most significant effect (**Fig. S1)**. We next conducted comparisons between sex for each measured protein in 1) sham-injured animals only (**Fig. 1B**) and 2) all sham and injured animals (**Fig. S1A**). Importantly, all proteins determined to be significantly different between sexes based on the sham animals alone were also significantly different between sexes when all samples were used, regardless of injury condition (**Fig. 1C**, **Fig. S2**, **Table S2**). An additional 12 and 17 significantly different proteins were identified in the frontal cortex and hippocampus, respectively, when all animals across injury conditions were compared (**Fig. 1C, Fig. S2**, **Table S2**).

Importantly, female samples showed significantly higher levels of Aβ42 in the frontal cortex compared to males, and higher total and phosphorylated tau at a pathologically relevant residue (T181) (74) in the hippocampus compared to male samples (p<0.0001, Wilcoxon rank-sum; **Fig. 1C**, **Table S2**). This matches most prior studies in 3xTg-AD mice, as reviewed in (72). Interestingly, we found significantly higher phospho-tau T181 in male compared to female frontal cortices (p<0.0001, Wilcoxon rank-sum). While our findings of sex differences are consistent with prior reports in 3xTg AD mice (72), there have been no reports evaluating sex differences in immune signaling or glial activation. Here, we found that the majority of the 46 measured immune and pathological marker proteins in both brain regions showed higher average levels in males compared to females (**Fig 1B**, **Fig S1A**). Of those, 31 from the frontal cortex (24 in sham-only) and 21 from the hippocampus (15 in sham-only) reached statistical significance (p<0.05, Wilcoxon rank-sum; **Fig. 1C**, **Fig. S2**, **Table S2**). Importantly, the astrocyte reactivity markers GFAP and S100B were significantly elevated in the male frontal cortex and hippocampus, indicating increased astrocyte reactivity in males. Taken together, these data suggest an elevated immune signaling baseline in young (2-4mo) male 3xTg-AD mice compared to their age-matched female counterparts.

To determine if differences in immune markers between male and female mice could be explained by the well-documented disparities in Aβ and tau in this model (72), we adjusted our data for effects related to Aβ (Aβ40 and Aβ42), tau (total tau and phospho-tau T181), or both (**Methods, Fig. S3, Table S3**). Adjustment for all pathological markers in the frontal cortex reduced the number of differentially expressed proteins from 31 to 15 elevated in males and from 9 to 3 elevated in females, suggesting that approximately half of all differentially expressed proteins can be explained by pathology. A similar effect is seen in the hippocampus after adjustment: from 21 to 12 proteins elevated in males and 16 to 8 in females. Interestingly, the removal of only Aβ-related effects (Aβ40 + Aβ42) in the hippocampus only accounted for 2 differentially expressed proteins (**Table S3**). The persistence of sexual dimorphism among immune markers (mostly cytokines) after adjustment for pathological measures suggests inherent differences in baseline immune signaling of young male and female 3xTg-AD mice independent of their stage of pathogenesis. Thus, we analyzed male and female data separately in this study.

### Cytokines and MAPK phospho-proteins are elevated after each closed-head injury

To identify patterns of co-varying cytokines and phospho-proteins with injury and time point, we next conducted hierarchical clustering within each sex and brain region (**Methods, Fig. S4-5**). Importantly, we found that females exhibited robust variation of these signaling proteins with both number of injuries and time point. The clustering identified a group of injury-induced cytokines (IL-9, IL-17, KC, IL-1α, IL-2, IL-1β, IL-13) and MAPK phospho-proteins (phospho-Atf2, phospho-Mek1) that tightly clustered together within the frontal cortex data from female mice (**Fig. S4, “Group 1”**). Proteins within this group were significantly increased at 24h after 1x, 2x, and 3xCHI, showed mild increases 24h after 4xCHI, and returned to baseline sham levels 24h after 5xCHI (**Fig. 2**), suggesting a dysregulated immune response after four or more injuries. Interestingly, these proteins were elevated 4h post-5xCHI compared to sham, 4h post-1xCHI, and 4h post-3xCHI animals (**Fig. 2**), indicating an earlier response after multiple injuries consistent with a “primed” immune state (75). Although many of the same cytokines and phospho-proteins from Group 1 again clustered together within the female hippocampus (**Fig. S4**) and male frontal cortex and hippocampus (**Fig. S5**), this same trend of an injury-dependent increase was only noted in the female frontal cortex. Together, these data indicate the importance of evaluating multiple time points after injury to fully define the immune response and emphasize the need to evaluate males and females separately.

**Figure 2.**
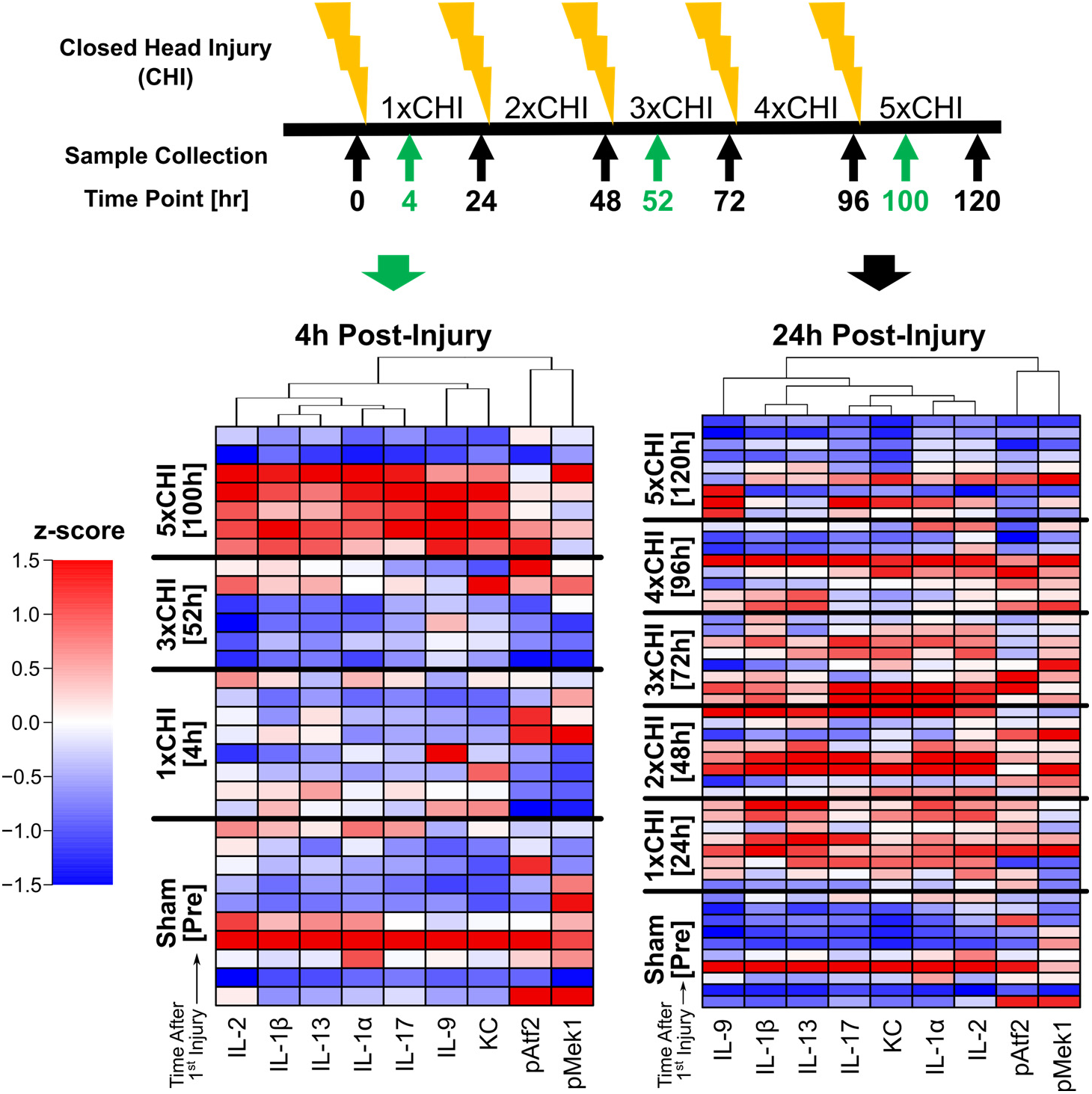
Cytokines and MAPK phospho-proteins are elevated after each closed-head injury in females. Hierarchical clustering of cytokines and MAPK phospho-proteins (subset from **Fig. S4, “Group 1”**) from the frontal cortex of female 2-4mo 3xTg sham-injured mice and 4hr (left) or 24hr (right) after 1-5x closed head injuries (CHI). Each row represents data from an individual mouse z-scored along each column.

### Protein and transcript markers of pathology are elevated after injury

Our motivation for conducting this study in 3xTg-AD mice was to identify relationships between immune signaling and molecular markers associated with AD-related pathology. In particular, TBI is known to cause neurodegenerative pathologies, such as neurofibrillary tangles associated with hyperphosphorylated tau as well as the build-up of Aβ plaques. To assess the relationship between our injury-elevated Group 1 immune signals and pathology-associated proteins, we quantified total tau, phospho-tau (T181), Aβ 1-40, and Aβ 1-42 by Luminex ELISA, then conducted clustering and correlation analyses. Few significant acute changes in Aβ or correlation with Group 1 proteins were observed in either region or sex (**Figs. S8-9**). Interestingly, however, in both brain regions and both sexes, the Group 1 immune markers clustered with phospho-tau with few exceptions (**Fig. S4-5**), suggesting a connection between cytokines and pathologically relevant tau phosphorylation (74). Pearson’s correlation coefficients between phospho-tau T181 and each cortical immune marker within each injury group (**Fig. 3A**) revealed that the majority of immune signals in Group 1 were correlated (|R|>0.50) across multiple time points. These data thus suggest that injury drives acute cytokine and MAPK immune signaling associated with increased phosphorylated tau.

**Figure 3.**
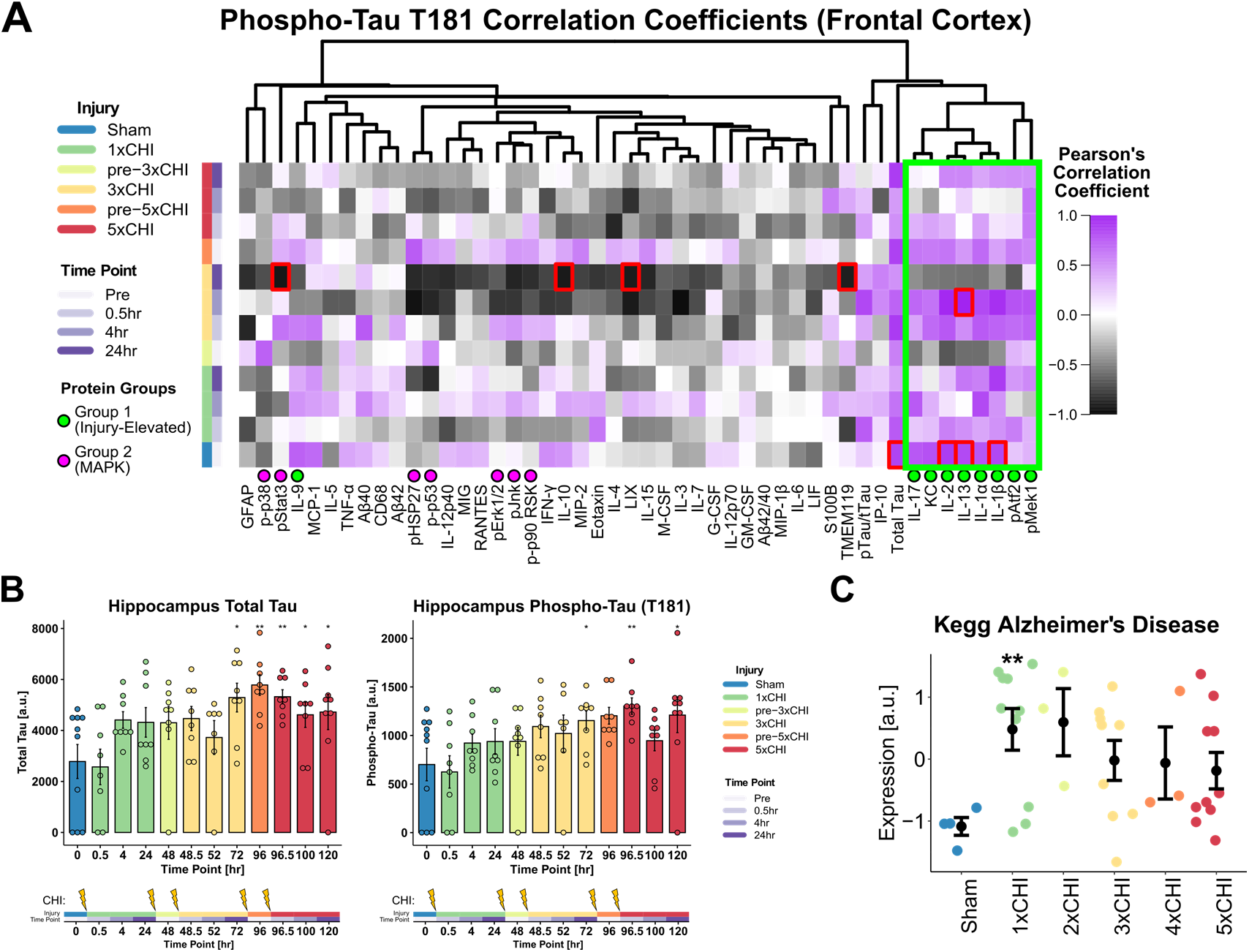
Total and phospho-tau increase with successive injuries in the hippocampus, AD-related genes increase in the somatomotor cortex after 1xCHI. **A**) Heatmap of Pearson’s correlation coefficient of cortical phospho-tau T181 versus each measured cortical protein (columns) within each injury group and time point (rows); red boxes indicate FDR-adjusted p<0.05; green and purple dots represent Group 1 and 2 proteins, respectively. Green box shows eight out of nine Group 1 proteins cluster together and show correlation coefficients R>0.50 with phospho-tau T181. **B**) Total tau is significantly upregulated in the hippocampus at 24hr post-3xCHI and each successive time point compared to sham-injured animals (*p<0.05, **p<0.01 Wilcox). Phospho-tau (T181) is significantly upregulated in the hippocampus 30min and 24hr post-5xCHI compared to sham-injured animals (*p<0.05, **p<0.01 Wilcox) (mean ± SEM). **C**) Expression of the gene set “Kegg Alzheimer’s Disease” significantly increases after 1xCHI (**p<0.01, t-test with Bonferroni correction) (mean ± SEM). See individual gene changes in **Fig. S10**.

Interestingly we observed these pronounced correlations in the frontal cortex despite only finding group-wise elevation of phospho-tau at 24h post-3xCHI in female frontal cortices (**Fig. S6**) and no group-wise changes in males. However, total tau and phospho-tau (T181) were elevated at multiple time points after 3x and 5xCHI (**Fig. 3B**). These data thus indicate that our CHI model induces acute pathological changes in a region-dependent manner, and that Group 1 immune signaling correlates with phosphorylated tau even in the absence of group changes in total or phosphorylated tau.

Due to the observed increases in total and phospho-tau in injured animals, we hypothesized that genes associated with AD pathogenesis may also become upregulated after rmTBI. To test this, we conducted bulk RNAseq on a subset of 40 females (somatomotor cortex, n=3-4 per group), then conducted gene set variation analysis (GSVA) using the Alzheimer’s disease-related gene set from the Molecular Signatures Database (MSigDB) C2 curated gene sets collection. Importantly, this gene set was significantly enriched after a single injury and remained enriched with subsequent injuries (**Fig. 3C, Fig. S10**). Thus, our protein and transcriptional data together indicate that pathological markers increase after injury and are correlated with neuroimmune markers.

Cytokines and MAPK phospho-proteins elevated after injury co-label with neurons Having found that Group 1 proteins correlated with phospho-tau, which is a primarily a neuronal protein (Fig. 3A), we next asked if Group 1 signaling would be associated with neurons or other cell types. To this end, we conducted immunohistochemistry for various Group 1 proteins together with markers for neurons (NeuN) and astrocytes (GFAP) at the 24h time point in mice exposed to 1x or 3xCHI. We found that IL-1*α*, IL-1*β*, IL-13, phospho-Atf2, and phospho-Mek1/2 co-localized with the neuronal marker NeuN in the cortex and hippocampus after injury, suggesting neuronal involvement in post-rmTBI immune signaling (Fig. 4, Fig. S11Error! Reference source not found. Figure). Additionally, IL-1β, IL-13, KC, phospho-Atf2, and phospho-Mek1/2 did not co-localize with the astrocyte marker GFAP in the cortex (**Fig. S12**) or hippocampus (**Fig. S13**).

**Figure 4.**
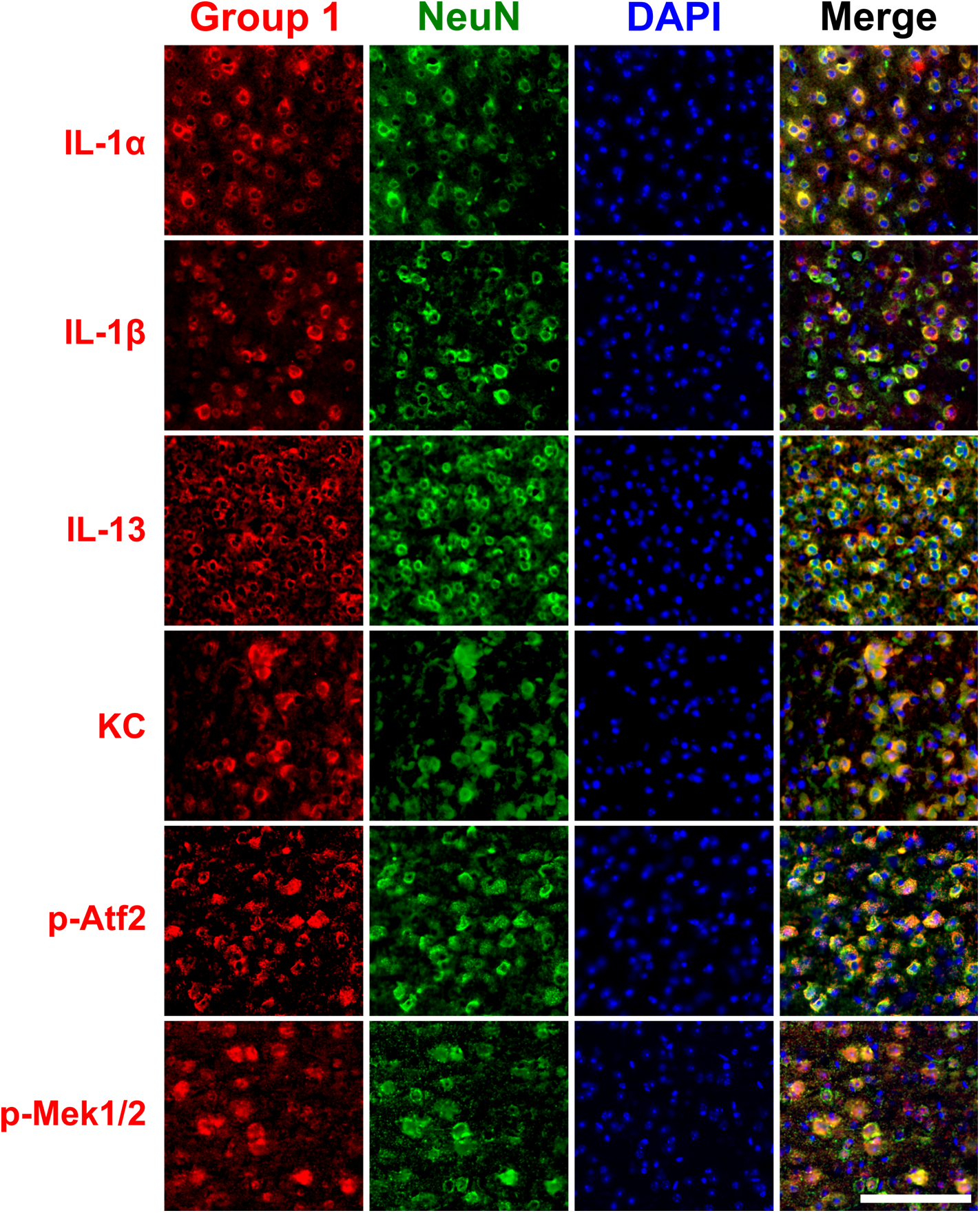
Group 1 cytokines and phospho-proteins co-label with NeuN in frontal cortex 24hr post-CHI. Group 1 cytokines (IL-1α, IL-1β, IL-13, and KC) and MAPK phospho-proteins (phospho-Atf2 and phospho-Mek1/2) (red) co-stained with NeuN (green) and DAPI (blue) show neuronal localization in the frontal cortex 24h post-CHI in female 3xTg mice aged 2-4mo (scale bar: 500μm).

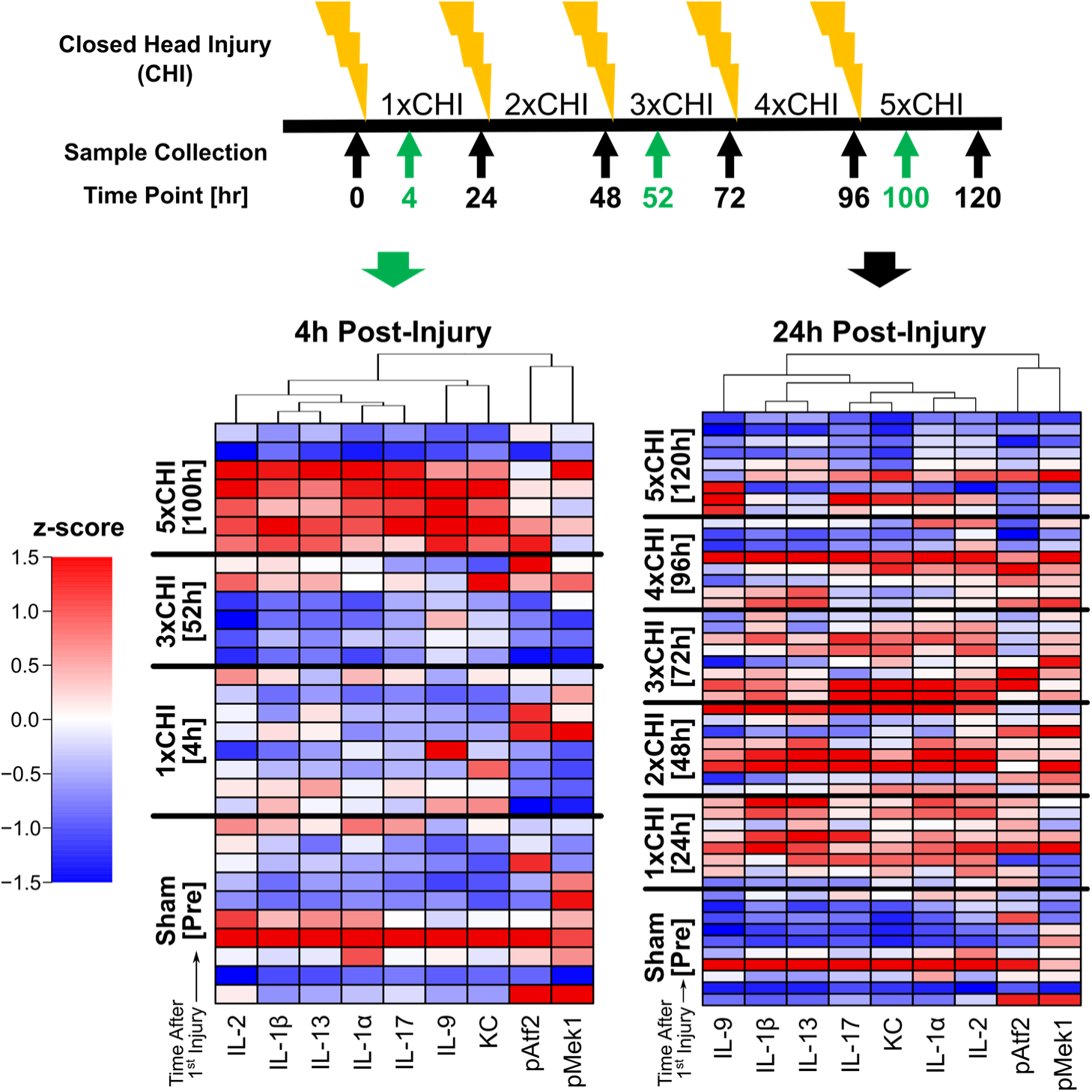

### Bulk RNAseq analysis reveals the effect of injury on pathology- and cell type-associated gene modules

To illuminate the broad effects of repetitive mTBI on cortical transcriptional networks, we next conducted network analysis on the same subset of 40 female cortical samples (somatomotor cortex, adjacent to the frontal cortex sections used for protein analysis) used in **Fig. 3C**, sampled from all injury groups and time points (n=3-4 per group). One sample was removed from the analysis due to its outlier status and exclusion within the protein data (Mahalanobis multivariate outlier detector with α<0.001, **Methods**).

To obtain an unbiased view of expression patterns within the data, we first conducted WGCNA (**Methods**), which identified 14 distinct modules of co-varying genes (**Fig. 5A**). A module eigengene (ME) was computed for each module as its first principal component, and expression scores of each ME were displayed for each sample across all injury and time point groups (**Fig. 5B**). Due to small sample sizes and relatively low variation within time points of each injury group (**Fig. 5B**), we combined post-injury time points (30m, 4h, 24h) into one group per injury number and compared module expression across number of injuries. The two modules comprised of the largest number of genes, ME1 (turquoise) and ME2 (blue), rapidly changed after 1xCHI, with ME1 showing a significant increase (p<0.05) and ME2 showing trending decreases (p=0.054) post-injury. In contrast, ME9 (magenta) progressively increased with successive injuries (**Fig. 5C**) and tracked closely with ME7 and ME8 (black, pink) after 2-5xCHI. Moreover, ME3 (brown) and ME6 (red) were increased after 1xCHI then decreased compared to sham after 5xCHI, while ME10 (purple) and ME11 (green-yellow) peaked after 2xCHI, then returned to sham levels. These modules therefore define changes associated with rapid (ME1, ME2), transient (ME3, ME6, ME10, ME11) and gradual time scales (ME7, 8, 9).

**Figure 5.**
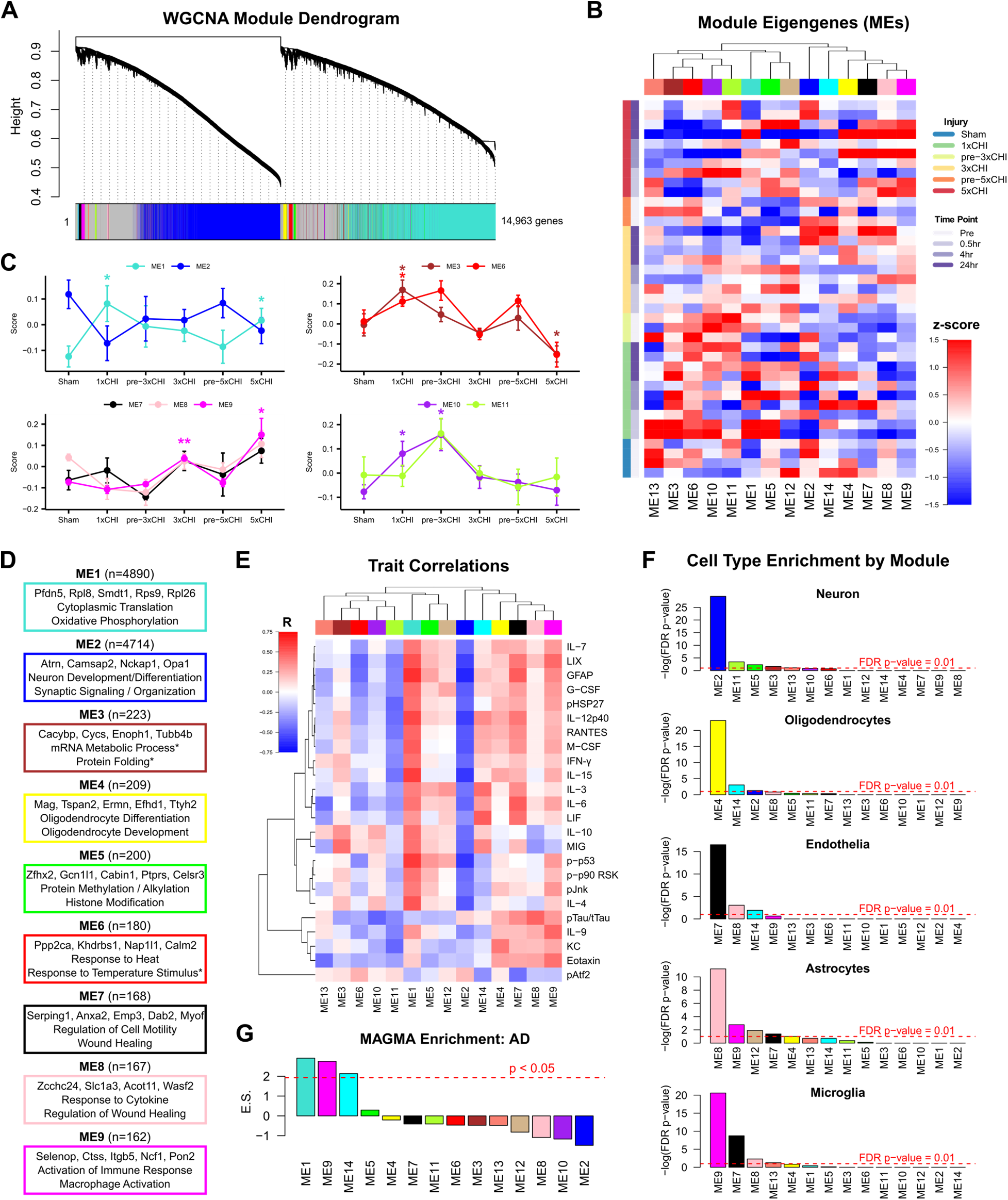
WGCNA of somatomotor cortex transcriptomes identifies modules of co-expressed genes associated with injury, GO biological processes, inflammatory protein expression, brain cell types, and AD genetic risk factors. **A**) WGCNA identified 14 modules of co-expressed genes. **B**) Expression scores for the 14 module eigengenes (MEs, the first principal component of each module) for each sample. MEs are clustered by Euclidean distance (columns) and expression scores are displayed by increasing injury number and time point from bottom to top of the heatmap (rows). **C**) Line plots show changing ME expression scores over increasing numbers of injuries. Time points for 30min, 4hr, and 24hr after each injury (1x, 3x, 5xCHI) are combined due to low sample sizes (n=2-4). ME1 (turquoise) and ME2 (blue) show opposite behavior after 1xCHI, with ME1 significantly increasing alongside trending decreases in ME2 after a single injury. ME 9 (magenta) shows significant increases after successive injuries that track closely with ME7 and 8 (black and pink) (*p<0.05, **p<0.01, permutation tests compared to sham, see **Methods**) (mean ± SEM). **D**) Gene ontology enrichment of biological processes was conducted for each module (Fisher’s exact test with FDR-corrected p-value <0.05). ME3 and ME6 had fewer than two significant GO sets. Sets marked with asterisks had FDR-corrected p>0.05 but uncorrected p<0.0001. **E**) Protein expression data from the frontal cortex of the same animals were correlated against each ME. Selected proteins showed a correlation p-value of at least 0.01 against at least one ME. Red indicates a positive Pearson’s correlation coefficient (R) while blue indicates negative correlation. **F**) Cell type enrichment analysis showed significant enrichment of neuronal genes in ME2 (blue), oligodendrocyte genes in ME4 (yellow), endothelia genes in ME7 (black), astrocyte genes in ME8 (pink), and microglial genes in ME9 (magenta). **G**) MAGMA enrichment for GWAS AD risk showed significant over-representation within ME1 (turquoise), ME9 (magenta), and ME14 (cyan). The dotted red line indicates an enrichment z-score of 1.96, above which a module is considered significantly enriched (normal distribution, p<0.05).

To understand the biological relevance of each module, we used gene ontology (GO, via PANTHER) to identify top GO terms enriched in each module (**Fig. 5D**). Importantly, the rapidly changing ME2 (blue) was enriched for genes related to nervous system development, suggesting a rapid downregulation of homeostatic neuronal functions. The gradually changing modules, ME7, ME8, and ME9 (black, pink, magenta), were enriched for GO terms associated with wound healing, cytokine response, gliogenesis, and regulation of the immune response, suggesting a multi-day response that involves immune processes and healing. Importantly, those modules that increase with injury were all correlated with immune proteins (inflammatory cytokines, MAPK activation, and GFAP) quantified in the frontal cortex of the same animals (ME1, 7, 9; “Trait Correlations” **Fig. 5E**), whereas ME2, which shows decreasing trends after injury and is associated with neuronal function, was inversely correlated with immune proteins (**Fig. 5E**).

We next conducted a cell-type enrichment analysis for genes with high specificity in neurons, oligodendrocytes, endothelia, astrocytes, and microglia derived from cell type specific mouse brain proteome and transcriptome studies Sharma et. al. (63) and Zhang et. al. (64) (**Fig. 5F**). We found that ME2 (blue), which decreases after a single injury and is inversely correlated with immune protein markers, was significantly enriched for neuronal genes. In contrast, ME7, 8, and 9 (black, pink, magenta), which show gradual increasing trends with successive injuries and (with the exception of ME8) correlate with immune protein markers, were enriched for endothelial, astrocyte, and microglial genes, respectively.

We concluded this unbiased analysis by asking if specific modules were associated with AD-related gene expression. To test this, we used a Multi-marker Analysis of GenoMic Annotation (MAGMA) analysis to determine if disease risk associated with AD-related genes was significantly elevated within specific gene modules. In this analysis, each module is evaluated for AD-enriched genes (previously identified by GWAS, 66–68) weighted by their associated risk factor for AD. Indeed, AD risk was significantly elevated in ME1 (i.e. *Pfdn5*, *Rps11*, *Rpl13a*, *Lamtor4*, *Rpl14*), which increases with a single injury and correlates significantly to several inflammatory proteins. AD risk was also elevated in ME9 (i.e. *Hepacam*, *Vav1*, *Hexb*, *Grn*, *Laptm5*), which increases gradually, correlates with inflammatory proteins, and is enriched for microglial genes (**Fig. 5G**). Taken together, these data suggest an early inflammatory response associated with AD genes (ME1) and a concomitant, trending, neuronal decrease in homeostatic functions (ME2) after a single injury. Moreover, multiple injuries progressively increased gene expression profiles associated with wound healing and regulation of inflammation within endothelia, astrocytes, and microglia.

### Transcriptional and protein markers of astrocyte reactivity are increased after repetitive closed head injury

Astrocyte reactivity is a central phenotype of brain injury. Several studies have reported elevated expression of the astrocyte reactivity marker GFAP within hours, and up to 1mo, after TBI. Moreover, GFAP is being evaluated as a CSF biomarker for TBI diagnosis and prognosis (76–80). We therefore hypothesized that astrocyte reactivity would be increased in 3xTg mice exposed to successive mTBIs. To analyze possible changes in astrocyte phenotypes and function following injury, we conducted a gene set variation analysis (GSVA) (56,69) of custom-annotated astrocyte gene sets that define astrocyte homeostatic and reactive functions (1,740 genes spanning 17 gene sets plus the A1 and A2 gene sets (81)) from our prior work (71) (**Fig. 6A**). To define the specific effects of injury and time point on astrocyte transcriptional profiles, we conducted multiple linear regression of the enrichment scores of all 17 astrocyte gene sets using number of injuries, time point after injury, and a binary injury variable (injured or sham) as independent variables. Consistent with our WGCNA, this analysis revealed that the number of injuries had a greater effect on gene set enrichment than the post-injury time point (**Fig. 6B**). We therefore combined post-injury time points (30min, 4h, 24h) to create a single group for each number of injuries 1-5xCHI, improving our ability to analyze differences despite our small sample sizes (n=3-4 per time point).

**Figure 6.**
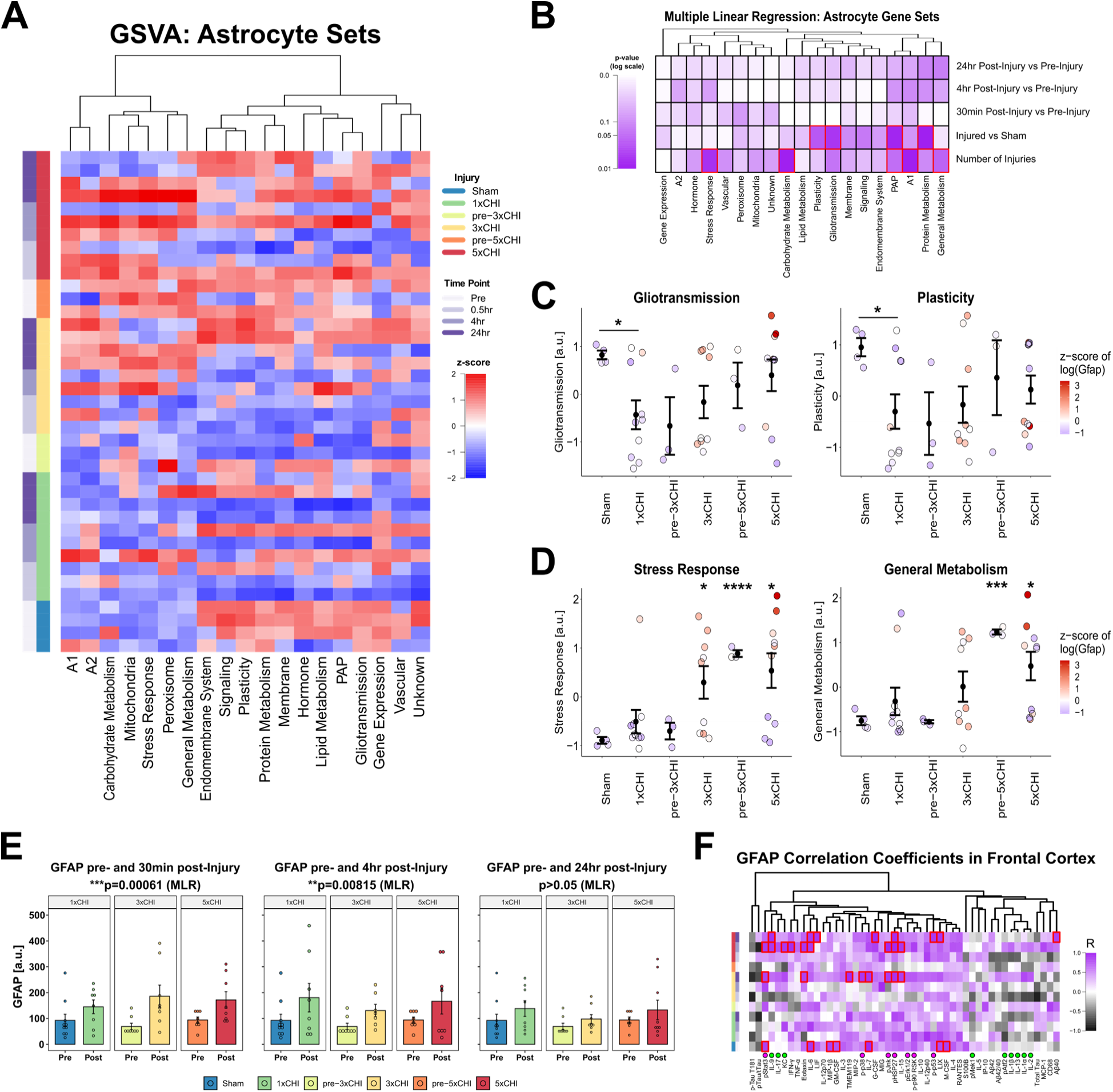
Cortical astrocytes become reactive following mild TBI. **A**) Gene set variation of custom-curated astrocyte gene sets (columns) across injury and time point groups (rows) reveals two distinct expression patterns: gradual increase by 3-5xCHI (left cluster) and rapid decrease after 1xCHI (right cluster). Gene sets are z-scored and clustered by Euclidean distance. **B**) Multiple linear regression models were calculated for each gene set using fixed effects of time point after most recent injury, binary injury status (injured, sham), and number of injuries. The model shows that the effects of injury (both numeric and binary) outweigh time-dependent changes for most gene sets. Red boxes indicate statistical significance (p<0.05). **C**) Astrocyte gene sets Gliotransmission and Plasticity are representative of right cluster, displaying a decrease after 1xCHI followed by return to sham levels by 4-5xCHI (*p<0.05, t-test with Bonferroni adjustment) (mean ± SEM). Data points are colored by the z-score of the log of astrocyte reactivity marker gene *Gfap*. **D**) Astrocyte gene sets Stress Response and A1 are representative of left cluster, displaying a gradual increase after 3-5xCHI (*p<0.05, ***p<0.001, ****p<0.0001; t-test with Bonferroni adjustment) (mean ± SEM). Data points are colored by the z-score of the log of astrocyte reactivity marker gene *Gfap*, which is associated with high expression in either set. **E**) Glial fibrillary acidic protein (GFAP) is significantly upregulated 30min and 4hr post-injury vs pre-injury (***p<0.001, **p<0.01; multiple linear regression model, pre/post and number of injuries) (mean ± SEM). **F**) Heatmap of Pearson’s correlation coefficient of cortical GFAP versus each measured cortical protein (columns) within each injury group and time point (rows); red boxes indicate FDR-corrected p<0.05. Panels A-D represent transcriptional data from female somatomotor cortex. Panels E-F represent protein data from female frontal cortex.

Interestingly, clustering of sample enrichment scores for the 17 astrocyte gene sets revealed two distinct patterns of expression after repeated injury (**Fig. 6A**). First, twelve astrocyte gene sets formed a cluster characterized by rapid decreases in enrichment following the first CHI, as represented by the Gliotransmission and Plasticity sets (**Fig. 6C**). These sets mostly showed recovery of expression to sham-levels by 24hr post-4/5xCHI. Many of these sets represent normal homeostatic astrocytic functions, such as gliotransmission, plasticity, gene expression, and the perisynaptic astrocyte process (PAP). A rapid decrease in expression of these sets following 1xCHI suggests astrocytes first respond to injury by altering their homeostatic functions. The second response pattern, represented by Stress Response and General Metabolism (**Fig. 6D**), shows a gradual increase in expression after 3-5xCHI. Coloring data points according to *Gfap* revealed an association between astrocyte reactivity and elevated Stress Response and General Metabolism gene sets. Other gene sets exhibiting this trend of a gradual increase include the A1 and A2 phenotypes and Carbohydrate Metabolism, which are functions that are generally related to astrocyte reactivity (81,82). These data therefore suggest elevated astrocyte reactivity by 3-5xCHI, 72-120hr after the first daily injury.

Since the RNAseq analysis suggested increased astrocyte reactivity at the transcriptional level, we next analyzed astrocyte reactivity at the protein level. To do so, we quantified astrocyte reactivity markers glial fibrillary acidic protein (GFAP) and S100 calcium binding protein B (S100B) by ELISA in the frontal cortex and hippocampus of all female mice exposed to 1x, 3x, or 5x once daily CHIs. To assess the behavior of GFAP in our injury model, we conducted a multiple linear regression analysis with dependent variables of time point following injury (pre-injury vs 30min, 4hr, or 24hr post-injury) and total number of injuries. Across female frontal cortex samples, we found that post-injury samples were significantly upregulated from pre-injury samples at both 30min and 4hr, but not 24hr post-injury (**Fig. 6E**), and that the total number of injuries was not a significant predictor of GFAP levels. In a parallel analysis, multiple linear regression of cortical S100B in female mice similarly revealed statistically significant upregulation 30min post-injury vs pre-injury but not after 4hr or 24hr (**Fig. S14**). Taken together, these data indicate a rapid, transient state of astrocyte reactivity following each repeated CHI that begins as early as 30min, but returns to basal levels by 24hr. Due to the significant increases after successive injuries in microglial-enriched and AD-GWAS-enriched ME9 (**Fig. 5**), we also quantified microglial activation marker CD68 but found that changes were minimal and inconsistent between sex and region (**Fig. S15**).

Because astrocyte reactivity is a key component of the brain’s response to stress and injury, we next analyzed the relationships between GFAP protein levels, cytokine expression, and MAPK signaling activity. Among both female cortical and hippocampal samples, a group of MAPK phospho-proteins, termed “Group 2” (phospho-Stat3, phospho-Jnk, phospho-HSP27, phospho-p38, phospho-p90 RSK, phospho-Erk1/2, and phospho-p53), clustered tightly together across all injury groups and closely with GFAP (**Fig. S4-5**), indicating potential involvement of astrocyte reactivity in both brain regions. Therefore, we next computed Pearson’s correlation coefficients between GFAP and each measured immune marker within each injury and time group (**Fig. 6F**) and found broad positive correlations both before and after injury, emphasizing the involvement of astrocytes in brain immune signaling.

## DISCUSSION

In this study, we comprehensively profiled immune and pathological responses to rmTBI in the brains of 3xTg-AD mice using a combination of targeted protein panels and RNA sequencing. We hypothesized that repeated mTBI drives progressive changes in immune signaling together with markers of astrocyte reactivity, microglial activation, and changes in tau and Aβ. Because wild type mice do not develop plaques and tangles associated with AD or brain injury-induced pathology, we employed the triple transgenic model of AD (3xTg-AD) in the present study to evaluate relationships between injury, pathological hallmarks, and markers of immune response across multiple acute time points. To our knowledge, this study is the first to define the temporal evolution of immune and pathological responses to rmTBI at both the protein and transcriptional levels in a mouse model capable of displaying both Aβ plaques and neurofibrillary tangles.

While sex differences in pathology and cognitive outcomes after injury in 3xTg mice have been extensively studied (72), few studies have evaluated sex differences in immune signaling. Indeed, to our knowledge all published brain injury studies which make use of the 3xTg-AD model report only male mice or combined-sex experimental groups. Sexual dimorphism of brain immune signaling in both humans and other mouse models is well-documented, generally showing a stronger immune response in females (83). Female 3x-Tg AD mice are reported to have increased Aβ pathology, possibly due to increased β-secretase processing (73,84,85) while male 3xTg AD mice and have been reported to have increased systemic autoimmunity (86), emphasizing the need to evaluate the role of sex. Indeed, we found that 40 out of 46 measured proteins were significantly different between males and females in the frontal cortex, as well as 37 out of 46 in the hippocampus. Further, we determined that total tau, phospho-tau T181, Aβ40, and Aβ42 accounted for only half of the differentially expressed proteins. In particular, pathology-adjusted data left 18 significantly different proteins in the frontal cortex and 20 in the hippocampus. These findings indicate that male and female samples should be analyzed separately. Indeed, dichotomizing our protein data into male and female datasets revealed that females exhibited a much stronger response to injury (**Fig. S4, Fig S5**). For this reason, we focused our analysis on females.

A key finding of our work is that many of the quantified cytokines and phospho-protein signaling molecules co-labeled with NeuN. Neurons are known to express a plethora of cytokine receptors and undergo phospho-protein signaling cascades (i.e., MAPK and NF-κB, among others) in response to inflammation (87–92), but an increasing body of work suggests that neurons also participate in cytokine secretion in both homeostatic and pathological conditions (93–104). Indeed, prior work from our own group has found increased MAPK and NF-κB signaling and expression of cytokines that co-label with neurons following rmTBI in male wild type mice (105). Moreover, multiple studies have reported evidence of co-localization of immune signaling proteins and/or their transcripts with neurons in most of our “Group 1” proteins, which are elevated in the cortex in females following each CHI: IL-1α (93,94), IL-1β (95,96), IL-2 (97,98), IL-9 (99), IL-13 (100–102), and KC. Importantly, our IHC in the current study revealed prominent neuronal co-localization of four cytokines (IL-1α, IL-1β, IL-13, and KC) and both MAPK phospho-proteins (phospho-Atf2 and phospho-Mek1) within “Group 1,” which is consistent with prior literature. Our data together with prior literature therefore support a role for neurons in orchestrating an immune signaling response post-rmTBI.

To our knowledge, our work is the first to conduct transcriptional profiling after successive mTBIs in a transgenic mouse model. We used WGCNA to provide an unbiased summary of gene expression trends in the data, then assessed each identified gene module for enhanced expression of specific biological processes (gene ontology), cell type specific markers, and GWAS-identified markers of AD, as well as correlation with paired protein expression (**Fig. 5**). Indeed, we identified an early inflammatory response associated with AD-GWAS gene expression and protein translation after a single injury, paired with a concomitant decrease in neuronal-enriched functions associated with neuronal development and synaptic signaling. Within days of repeated injury, there was a gradual increase of endothelia-, astrocyte-, and microglia-enriched gene modules associated with wound healing and regulation of inflammation. Interestingly, we also found a significant enrichment of AD-GWAS risk genes within the microglia-enriched gene modules, suggesting microglial involvement in acute pathogenesis after injury. Collectively, these data suggest that even a single mTBI elicits a change in genes associated with lost neuronal homeostasis, whereas repeated daily mTBIs yield changes that propagate into astrocytes, microglia, and endothelial cells.

Interestingly, our protein data showed that the astrocyte reactivity markers GFAP and S100B are transiently elevated at 30min after each repetitive mTBI in the female frontal cortex. Although few studies have characterized such rapid changes in reactivity markers following closed-head injury, elevated GFAP and GFAP breakdown products are detectable in serum less than one hour after mild-to-severe TBI in mice (76–78). This parallels findings in human patients, where serum GFAP levels rapidly increase within an hour then decline within 24-72hr following mild-to-severe injury (78–80,106–108) as well as some mouse studies (109,110), but is in contrast to several other studies in rodents showing sustained GFAP elevation up to two weeks post-injury (111–114). Differences in rodent and injury models, sample type (serum, CSF, tissue lysate), and sex composition of experimental groups may account for some of these conflicting results. On the transcriptional level, we found that astrocyte-specific gene sets for stress response, general metabolism, and immune-related phenotypes A1 and A2 (81) showed gradual increases compared to sham after 3-5xCHI, while homeostatic sets associated with gliotransmission and astrocyte plasticity showed decreased expression compared to sham after just 1xCHI. These findings mirror previous studies which have identified elevated immune response genes in astrocytes 7 days post-severe TBI (115) and the lack of astrocyte-specific inflammatory gene changes 24h after mild TBI (116). Combined, our protein and transcriptional data indicate that astrocytes are responsive to mTBI as early as 30min after injury, while transcriptional changes in stress response pathways persist on the order of days.

The MAPK intracellular signaling pathway is a known mediator of astrocyte reactivity (117–119), including after a traumatic brain injury (35). The clustering of female cortical GFAP and multiple MAPK phospho-proteins suggests that MAPK signaling may be responsible for the phenotypic transition to a reactive state in our model. Further, GFAP also correlates with a host of cytokines such as LIX, IL-7, IL-3, IL-12p40, RANTES, IL-15, M-CSF, G-CSF. These relationships generally exist independently of injury and suggest involvement within reactive astrocyte signaling, which peaks on a time scale of minutes-to-hours after each injury. Further studies should investigate the effect of MAPK signaling in astrocyte response to brain injury.

The current study has several limitations requiring future work. First, while the 3xTg-AD mouse model is becoming relatively popular for brain injury research (42–49), its predisposition towards spontaneous development of pathology due to the expression of three causal neurodegenerative transgenes is not genetically representative of most human TBI patients. The use of human transgenes is necessary due to the lack of pathogenicity of native mouse tau and Aβ, while the use of mutations specifically associated with dementia is justified by the short life span of mice. Here, we mitigated the potential effects of pre-existing pathology on injury outcomes by using mice aged to 2-4 months, well before the reported build-up of Aβ and phosphorylated tau at 6 and 12 months, respectively (120,121). Second, while bulk tissue processing enabled appropriate sample sizes and comparison between 12 injury and time point groups, it also limited our ability to draw conclusions at the level of a single cell. Our bulk analysis highlights injury groups and time points of interest to be interrogated in future work for single cell analysis. Single cell profiling would identify how subtypes of neurons, glia, and other cells respond to successive mTBIs at a higher resolution than can be inferred by network analysis of bulk transcriptomic data. Specifically, single cell analysis may help clarify the extent to which neurons may be involved in immune signaling through cytokine expression and MAPK activation as seen in the injury-driven “Group 1”. Third, our findings are purely correlative. While we have established evidence of a relationship between elevated neuroimmune signaling and outcome, future studies are required to establish whether this signaling indeed drives pathogenesis after rmTBI, or is merely correlated.

## CONCLUSIONS

In total, our data define an acute neuroimmune cascade of mild traumatic brain injury in 3xTg-AD mice, consisting of i) an immediate decrease in neuronal homeostatic gene expression and an elevation of AD-associated genes after a single injury, ii) elevation of a subset of cortical cytokines that co-label with neurons and correlated with phosphorylated tau, and iii) increased expression of non-neuronal genes suggesting glial reactivity within days of repeated injury. We provide resolution of these acute protein and transcriptional changes at a previously uncharacterized minutes-to-days time scale alongside the key experimental variables of sex, brain region, and number of injuries. Pronounced changes in neuronal gene expression that correlate with inflammatory protein expression and co-labeling of injury-elevated cytokines and MAPK phospho-proteins with neuronal marker NeuN suggest a key role for neurons in the modulation of neuroimmune activity following brain injury. Further, the association of phospho-Mek1 and phospho-Atf2, cytokines, and phospho-tau T181 after injury may represent a therapeutic avenue for regulating inflammation and acute pathological mechanisms. Future work should explore the causal functions of these key molecular signals.

## DATA AVAILABILITY

The gene expression FASTQ files and count matrix that support the findings of this study have been deposited in the Gene Expression Omnibus (GEO) repository under series record GSE226838. Protein expression data and mouse metadata has been deposited in the Open Data Commons for Traumatic Brain Injury repository.

## ABBREVIATIONS

3xTg-AD: Triple transgenic Alzheimer’s disease mouse model
Aβ: Amyloid beta
ACURO: Animal Care and Use Review Office (U.S. Army MRDC)
AD: Alzheimer’s disease
APP: Amyloid precursor protein
ARRIVE: Animal Research: Reporting of *In Vivo* Experiments
ATP: Adenosine triphosphate
BSA: Bovine serum albuminCD68: Cluster of differentiation 68
CHI: Closed-head injury
CRAN: Comprehensive R Archive Network
DAPI: 4′,6-diamidino-2-phenylindole
ELISA: Enzyme-linked immunosorbent assay
FDR: False discovery rate
G-CSF: Granulocyte colony-stimulating factor
GEO: Gene expression omnibus
GFAP: Glial fibrillary acidic protein
GM-CSF: Granulocyte macrophage colony-stimulating factor
GSVA: Gene set variation analysis
GWAS: Genome-wide association study
IACUC: Institutional Animal Care and Use Committee
Iba1: Ionized calcium binding adaptor molecule 1
IFN-γ: Interferon gamma
IHC: Immunohistochemistry
IL: Interleukin
IP-10: Interferon gamma-induced protein 10
KC: Keratinocyte chemoattractant
LIF: Leukemia inhibitory factor
LIX: Lipopolysaccharide-induced CXC chemokine
MAGMA: Multi-marker Analysis of GenoMic Annotation
MAPK: Mitogen-activated protein kinase
MAPT: Microtubule-associated protein tau
MCP-1: Monocyte chemoattractant protein 1
M-CSF: Macrophage colony-stimulating factor
MIG: Monokine induced by interferon gamma
MIP: Macrophage inflammatory protein
MRDC: Medical Research and Development Command (U.S. Army)
NCBI: National Center for Biotechnology Information
mTBI: Mild traumatic brain injury
PAP: Perisynaptic astrocyte processes
PCA: Principal component analysis
PD: Parkinson’s disease
PSEN1: Presenilin 1
pTau: Phospho-tau
RANTES: Regulated upon activation, normal T cell expressed and presumably secreted
rmTBI: Repetitive traumatic brain injury
RNA: Ribonucleic acid
S100B: S100 calcium binding protein B
TBI: Traumatic brain injury
TBST: Tris-buffered saline with tween
TMEM119: Transmembrane protein 119
TNF-α: Tumor necrosis factor α
tTau: Total tau
WGCNA: Weighted gene co-expression network analysis

## ACKNOWLEDGEMENTS

We wish to acknowledge fruitful discussions with Michelle LaPlaca, and Manu Platt. We also thank the core facilities at the Parker H. Petit Institute for Bioengineering and Bioscience at the Georgia Institute of Technology for use of their shared equipment, services, and expertise.

## FUNDING

This work was supported by the U.S. Department of Defense through the Congressionally Directed Medical Research Programs (CDMRP) under Award No. W81XWH-18-1-0669 (LBW/EMB). Opinions, interpretations, conclusions, and recommendations are those of the author and are not necessarily endorsed by the Department of Defense. It was also supported by the National Institutes of Health under Award Nos. 1R01NS115994 (LBW/EB), 1R01AG075820 (SR/LBW), 1R01NS114130 (SR), 1RF1AG071587 (SR). AFP was supported in part by the National Institutes of Health Cell and Tissue Engineering Biotechnology Training Grant (T32-GM008433).

## AUTHOR INFORMATION

### Contributions

AFP conducted data analysis, prepared all figures, and wrote the manuscript. ROB and AR conducted animal experiments, curated animal data, and conducted cerebral blood flow measurements. AFP, SB, AK, FRM, KU, and BD conducted molecular assays and experimentation. ED and SR contributed to informatic analysis. LBW and EMB conceived the study, supervised research, and revised the manuscript. All authors reviewed the manuscript.

### Corresponding Authors

Correspondence to Levi B. Wood and Erin M. Buckley.

## ETHICS DECLARATIONS

### Ethics approval and consent to participate

All protocols were approved by the U.S. Army Medical Research and Development Command (MRDC) Animal Care and Use Review Office (ACURO) and the Emory University Institutional Animal Care and Use Committee (IACUC) in accordance with National Institutes of Health guidelines.

### Consent for publication

Not applicable.

### Competing interests

The authors have no conflicts of interest to disclose.

## SUPPLEMENTARY INFORMATION

### Additional file 1

Supplementary Figures.

## Notes

### Competing Interest Statement

The authors have declared no competing interest.

## REFERENCES

1. US Department of Health & Human Services; Centers for Disease Control (CDC); National Center for Injury Prevention and Control. Report to Congress on Mild Traumatic Brain Injury in the United States: Steps to Prevent a Serious Public Health Problem: (371602004-001) [Internet]. American Psychological Association; 2003 [cited 2021 Jan 9]. Available from: http://doi.apa.org/get-pe-doi.cfm?doi=10.1037/e371602004-001

2. Surveillance Report of Traumatic Brain Injury-related Emergency Department Visits, Hospitalizations, and Deaths. :24.

3. Iraji A, Benson RR, Welch RD, O’Neil BJ, Woodard JL, Ayaz SI, et al. Resting State Functional Connectivity in Mild Traumatic Brain Injury at the Acute Stage: Independent Component and Seed-Based Analyses. J Neurotrauma. 2015 07-15;32(14):1031–45.

4. McCrory P, Meeuwisse W, Aubry M, Cantu B, Dvorak J, Echemendia RJ, et al. Consensus statement on concussion in sport–the 4th International Conference on Concussion in Sport held in Zurich, November 2012. Clin J Sport Med. 2013 03;23(2):89–117.

5. Belanger HG, Vanderploeg RD, Curtiss G, Warden DL. Recent neuroimaging techniques in mild traumatic brain injury. J Neuropsychiatry Clin Neurosci. 2007;19(1):5–20.

6. Sours C, Zhuo J, Roys S, Shanmuganathan K, Gullapalli RP. Disruptions in Resting State Functional Connectivity and Cerebral Blood Flow in Mild Traumatic Brain Injury Patients. PLoS ONE. 2015;10(8):e0134019.

7. Guskiewicz KM, McCrea M, Marshall SW, Cantu RC, Randolph C, Barr W, et al. Cumulative effects associated with recurrent concussion in collegiate football players: the NCAA Concussion Study. Jama. 2003 11-19;290(19):2549–55.

8. Longhi L, Saatman KE, Fujimoto S, Raghupathi R, Meaney DF, Davis J, et al. Temporal window of vulnerability to repetitive experimental concussive brain injury. Neurosurgery. 2005 Feb;56(2):364–74; discussion 364-374.

9. Mez J, Daneshvar DH, Kiernan PT, Abdolmohammadi B, Alvarez VE, Huber BR, et al. Clinicopathological Evaluation of Chronic Traumatic Encephalopathy in Players of American Football. JAMA. 2017 Jul 25;318(4):360–70.

10. Fleminger S, Oliver DL, Lovestone S, Rabe-Hesketh S, Giora A. Head injury as a risk factor for Alzheimer’s disease: the evidence 10 years on; a partial replication. J Neurol Neurosurg Psychiatry. 2003 Jul;74(7):857–62.

11. Krueger F, Pardini M, Huey ED, Raymont V, Solomon J, Lipsky RH, et al. The role of the Met66 brain-derived neurotrophic factor allele in the recovery of executive functioning after combat-related traumatic brain injury. J Neurosci Off J Soc Neurosci. 2011 Jan 12;31(2):598– 606.

12. Heneka MT, Carson MJ, El Khoury J, Landreth GE, Brosseron F, Feinstein DL, et al. Neuroinflammation in Alzheimer’s Disease. Lancet Neurol. 2015 Apr;14(4):388–405.

13. Bower JH, Maraganore DM, Peterson BJ, McDonnell SK, Ahlskog JE, Rocca WA. Head trauma preceding PD: a case-control study. Neurology. 2003 May 27;60(10):1610–5.

14. Uryu K, Giasson BI, Longhi L, Martinez D, Murray I, Conte V, et al. Age-dependent synuclein pathology following traumatic brain injury in mice. Exp Neurol. 2003 Nov;184(1):214–24.

15. Tansey MG, Goldberg MS. Neuroinflammation in Parkinson’s disease: its role in neuronal death and implications for therapeutic intervention. Neurobiol Dis. 2010 Mar;37(3):510–8.

16. Simon DW, McGeachy MJ, Bayır H, Clark RSB, Loane DJ, Kochanek PM. The far-reaching scope of neuroinflammation after traumatic brain injury. Nat Rev Neurol. 2017 Mar;13(3):171–91.

17. Schimmel SJ, Acosta S, Lozano D. Neuroinflammation in traumatic brain injury: A chronic response to an acute injury. Brain Circ. 2017;3(3):135–42.

18. Morganti-Kossmann MC, Satgunaseelan L, Bye N, Kossmann T. Modulation of immune response by head injury. Injury. 2007 Dec;38(12):1392–400.

19. Loane DJ, Kumar A, Stoica BA, Cabatbat R, Faden AI. Progressive neurodegeneration after experimental brain trauma: association with chronic microglial activation. J Neuropathol Exp Neurol. 2014 Jan;73(1):14–29.

20. Bermpohl D, You Z, Lo EH, Kim HH, Whalen MJ. TNF alpha and Fas mediate tissue damage and functional outcome after traumatic brain injury in mice. J Cereb Blood Flow Metab Off J Int Soc Cereb Blood Flow Metab. 2007 11;27(11):1806–18.

21. Ebert SE, Jensen P, Ozenne B, Armand S, Svarer C, Stenbaek DS, et al. Molecular imaging of neuroinflammation in patients after mild traumatic brain injury: a longitudinal 123I-CLINDE single photon emission computed tomography study. Eur J Neurol. 2019;26(12):1426–32.

22. Perez-Polo JR, Rea HC, Johnson KM, Parsley MA, Unabia GC, Xu G, et al. Inflammatory consequences in a rodent model of mild traumatic brain injury. J Neurotrauma. 2013 May 1;30(9):727–40.

23. Mountney A, Boutté AM, Cartagena CM, Flerlage WF, Johnson WD, Rho C, et al. Functional and Molecular Correlates after Single and Repeated Rat Closed-Head Concussion: Indices of Vulnerability after Brain Injury. J Neurotrauma. 2017 Oct 1;34(19):2768–89.

24. Vállez García D, Otte A, Dierckx RAJO, Doorduin J. Three Month Follow-Up of Rat Mild Traumatic Brain Injury: A Combined [18F]FDG and [11C]PK11195 Positron Emission Study. J Neurotrauma. 2016 Oct 15;33(20):1855–65.

25. Rathbone ATL, Tharmaradinam S, Jiang S, Rathbone MP, Kumbhare DA. A review of the neuro- and systemic inflammatory responses in post concussion symptoms: Introduction of the “post-inflammatory brain syndrome” PIBS. Brain Behav Immun. 2015 May;46:1–16.

26. Giza CC, Hovda DA. The Neurometabolic Cascade of Concussion. J Athl Train. 2001 Sep;36(3):228–35.

27. Giza CC, Hovda DA. The New Neurometabolic Cascade of Concussion. Neurosurgery. 2014 Oct;75(0 4):S24–33.

28. Burda JE, Bernstein AM, Sofroniew MV. Astrocyte roles in traumatic brain injury. Exp Neurol. 2016 Jan;275(0 3):305–15.

29. Loane DJ, Kumar A. Microglia in the TBI Brain: The Good, The Bad, And The Dysregulated. Exp Neurol. 2016 01;275(0 3):316–27.

30. Witcher KG, Bray CE, Chunchai T, Zhao F, O’Neil SM, Gordillo AJ, et al. Traumatic Brain Injury Causes Chronic Cortical Inflammation and Neuronal Dysfunction Mediated by Microglia. J Neurosci. 2021 Feb 17;41(7):1597–616.

31. Kumar A, Loane DJ. Neuroinflammation after traumatic brain injury: Opportunities for therapeutic intervention. Brain Behav Immun. 2012 Nov 1;26(8):1191–201.

32. Ahmed SM, Rzigalinski BA, Willoughby KA, Sitterding HA, Ellis EF. Stretch-induced injury alters mitochondrial membrane potential and cellular ATP in cultured astrocytes and neurons. J Neurochem. 2000 May;74(5):1951–60.

33. Verderio C, Matteoli M. ATP mediates calcium signaling between astrocytes and microglial cells: modulation by IFN-gamma. J Immunol Baltim Md 1950. 2001 May 15;166(10):6383–91.

34. Neary JT, Kang Y, Willoughby KA, Ellis EF. Activation of Extracellular Signal-Regulated Kinase by Stretch-Induced Injury in Astrocytes Involves Extracellular ATP and P2 Purinergic Receptors. J Neurosci. 2003 Mar 15;23(6):2348–56.

35. Huang T, Solano J, He D, Loutfi M, Dietrich WD, Kuluz JW. Traumatic Injury Activates MAP Kinases in Astrocytes: Mechanisms of Hypothermia and Hyperthermia. J Neurotrauma. 2009 Sep;26(9):1535–45.

36. Neary JT, Kang Y, Tran M, Feld J. Traumatic injury activates protein kinase B/Akt in cultured astrocytes: role of extracellular ATP and P2 purinergic receptors. J Neurotrauma. 2005 Apr;22(4):491–500.

37. Floyd CL, Gorin FA, Lyeth BG. Mechanical strain injury increases intracellular sodium and reverses Na+/Ca2+ exchange in cortical astrocytes. Glia. 2005 Jul;51(1):35–46.

38. Bowman CL, Ding JP, Sachs F, Sokabe M. Mechanotransducing ion channels in astrocytes. Brain Res. 1992 Jul 3;584(1–2):272–86.

39. Ojo JO, Mouzon BC, Crawford F. Repetitive head trauma, chronic traumatic encephalopathy and tau: Challenges in translating from mice to men. Exp Neurol. 2016 Jan 1;275:389–404.

40. Bolton-Hall AN, Hubbard WB, Saatman KE. Experimental Designs for Repeated Mild Traumatic Brain Injury: Challenges and Considerations. J Neurotrauma. 2019 Apr 15;36(8):1203–21.

41. McAteer KM, Turner RJ, Corrigan F. Animal models of chronic traumatic encephalopathy. Concussion [Internet]. 2017 May 19 [cited 2021 Jan 12];2(2). Available from: https://www.ncbi.nlm.nih.gov/pmc/articles/PMC6093772/

42. Washington PM, Morffy N, Parsadanian M, Zapple DN, Burns MP. Experimental traumatic brain injury induces rapid aggregation and oligomerization of amyloid-beta in an Alzheimer’s disease mouse model. J Neurotrauma. 2014 Jan 1;31(1):125–34.

43. Tran HT, LaFerla FM, Holtzman DM, Brody DL. Controlled Cortical Impact Traumatic Brain Injury in 3xTg-AD Mice Causes Acute Intra-Axonal Amyloid-β Accumulation and Independently Accelerates the Development of Tau Abnormalities. J Neurosci. 2011 Jun 29;31(26):9513–25.

44. Winston CN, Noël A, Neustadtl A, Parsadanian M, Barton DJ, Chellappa D, et al. Dendritic Spine Loss and Chronic White Matter Inflammation in a Mouse Model of Highly Repetitive Head Trauma. Am J Pathol. 2016 Mar 1;186(3):552–67.

45. Tran HT, Sanchez L, Esparza TJ, Brody DL. Distinct Temporal and Anatomical Distributions of Amyloid-β and Tau Abnormalities following Controlled Cortical Impact in Transgenic Mice. PLOS ONE. 2011 Sep 29;6(9):e25475.

46. Saber M, Murphy SM, Cho Y, Lifshitz J, Rowe RK. Experimental diffuse brain injury and a model of Alzheimer’s disease exhibit disease-specific changes in sleep and incongruous peripheral inflammation. J Neurosci Res. 2021;99(4):1136–60.

47. Wu Z, Wang ZH, Liu X, Zhang Z, Gu X, Yu SP, et al. Traumatic brain injury triggers APP and Tau cleavage by delta-secretase, mediating Alzheimer’s disease pathology. Prog Neurobiol. 2020;185.

48. Bennett RE, Esparza TJ, Lewis HA, Kim E, Mac Donald CL, Sullivan PM, et al. Human apolipoprotein E4 worsens acute axonal pathology but not amyloid-β immunoreactivity after traumatic brain injury in 3xTG-AD mice. J Neuropathol Exp Neurol. 2013;72(5):396–403.

49. Hu W, Tung YC, Zhang Y, Liu F, Iqbal K. Involvement of activation of asparaginyl endopeptidase in tau hyperphosphorylation in repetitive mild traumatic brain injury. J Alzheimers Dis. 2018;64(3):709–22.

50. Meehan WP 3rd, Zhang J, Mannix R, Whalen MJ. Increasing recovery time between injuries improves cognitive outcome after repetitive mild concussive brain injuries in mice. Neurosurgery. 2012 Oct;71(4):885–91.

51. Brothers RO, Bitarafan S, Pybus AF, Wood LB, Buckley EM. Systems Analysis of the Neuroinflammatory and Hemodynamic Response to Traumatic Brain Injury. J Vis Exp JoVE. 2022 May 27;(183).

52. Wickham H, Averick M, Bryan J, Chang W, McGowan LD, François R, et al. Welcome to the Tidyverse. J Open Source Softw. 2019 Nov 21;4(43):1686.

53. Zhao S, Guo Y, Sheng Q, Shyr Y. Advanced heat map and clustering analysis using heatmap3. BioMed Res Int. 2014;2014:986048.

54. Wickham H, Chang W, Henry L, Pedersen TL, Takahashi K, Wilke C, et al. ggplot2: Create Elegant Data Visualisations Using the Grammar of Graphics [Internet]. 2022 [cited 2022 Oct 7]. Available from: https://CRAN.R-project.org/package=ggplot2

55. Kassambara A. ggpubr: “ggplot2” Based Publication Ready Plots [Internet]. 2020 [cited 2022 Oct 7]. Available from: https://CRAN.R-project.org/package=ggpubr

56. Hänzelmann S, Castelo R, Guinney J. GSVA: gene set variation analysis for microarray and RNA-seq data. BMC Bioinformatics. 2013 Jan 16;14:7.

57. 57. Coombes KR. Object Oriented Microarray and Proteomics Analysis (OOMPA). :6.

58. Love MI, Huber W, Anders S. Moderated estimation of fold change and dispersion for RNA-seq data with DESeq2. Genome Biol. 2014 Dec 5;15(12):550.

59. Langfelder P, Horvath S. WGCNA: an R package for weighted correlation network analysis. BMC Bioinformatics. 2008 Dec;9(1):1–13.

60. Johnson ECB, Carter EK, Dammer EB, Duong DM, Gerasimov ES, Liu Y, et al. Large-scale deep multi-layer analysis of Alzheimer’s disease brain reveals strong proteomic disease-related changes not observed at the RNA level. Nat Neurosci. 2022 Feb;25(2):213–25.

61. Johnson ECB, Dammer EB, Duong DM, Ping L, Zhou M, Yin L, et al. Large-scale proteomic analysis of Alzheimer’s disease brain and cerebrospinal fluid reveals early changes in energy metabolism associated with microglia and astrocyte activation. Nat Med. 2020 May;26(5):769–80.

62. Seyfried NT, Dammer EB, Swarup V, Nandakumar D, Duong DM, Yin L, et al. A Multi-network Approach Identifies Protein-Specific Co-expression in Asymptomatic and Symptomatic Alzheimer’s Disease. Cell Syst. 2017 Jan 25;4(1):60–72.e4.

63. Sharma K, Schmitt S, Bergner CG, Tyanova S, Kannaiyan N, Manrique-Hoyos N, et al. Cell type- and brain region-resolved mouse brain proteome. Nat Neurosci. 2015 Dec;18(12):1819– 31.

64. Zhang Y, Chen K, Sloan SA, Bennett ML, Scholze AR, O’Keeffe S, et al. An RNA-Sequencing Transcriptome and Splicing Database of Glia, Neurons, and Vascular Cells of the Cerebral Cortex. J Neurosci. 2014 09-03;34(36):11929–47.

65. Leeuw CA de, Mooij JM, Heskes T, Posthuma D. MAGMA: Generalized Gene-Set Analysis of GWAS Data. PLOS Comput Biol. 2015 Apr 17;11(4):e1004219.

66. Kunkle BW, Grenier-Boley B, Sims R, Bis JC, Damotte V, Naj AC, et al. Genetic meta-analysis of diagnosed Alzheimer’s disease identifies new risk loci and implicates Aβ, tau, immunity and lipid processing. Nat Genet. 2019 Mar;51(3):414–30.

67. Lambert JC, Ibrahim-Verbaas CA, Harold D, Naj AC, Sims R, Bellenguez C, et al. Meta-analysis of 74,046 individuals identifies 11 new susceptibility loci for Alzheimer’s disease. Nat Genet. 2013 Dec;45(12):1452–8.

68. Bellenguez C, Küçükali F, Jansen IE, Kleineidam L, Moreno-Grau S, Amin N, et al. New insights into the genetic etiology of Alzheimer’s disease and related dementias. Nat Genet. 2022 Apr;54(4):412–36.

69. Subramanian A, Tamayo P, Mootha VK, Mukherjee S, Ebert BL, Gillette MA, et al. Gene set enrichment analysis: a knowledge-based approach for interpreting genome-wide expression profiles. Proc Natl Acad Sci U S A. 2005 10-25;102(43):15545–50.

70. Liberzon A, Subramanian A, Pinchback R, Thorvaldsdóttir H, Tamayo P, Mesirov JP. Molecular signatures database (MSigDB) 3.0. Bioinformatics. 2011 Jun 15;27(12):1739–40.

71. Galea E, Weinstock LD, Larramona-Arcas R, Pybus AF, Giménez-Llort L, Escartin C, et al. Multi-transcriptomic analysis points to early organelle dysfunction in human astrocytes in Alzheimer’s disease. Neurobiol Dis. 2022 May;166:105655.

72. Dennison JL, Ricciardi NR, Lohse I, Volmar CH, Wahlestedt C. Sexual Dimorphism in the 3xTg-AD Mouse Model and Its Impact on Pre-Clinical Research. J Alzheimers Dis. 2021 Jan 1;80(1):41–52.

73. Hirata-Fukae C, Li HF, Hoe HS, Gray AJ, Minami SS, Hamada K, et al. Females exhibit more extensive amyloid, but not tau, pathology in an Alzheimer transgenic model. Brain Res. 2008 Jun 24;1216:92–103.

74. Goetzl EJ, Elahi FM, Mustapic M, Kapogiannis D, Pryhoda M, Gilmore A, et al. Altered levels of plasma neuron-derived exosomes and their cargo proteins characterize acute and chronic mild traumatic brain injury. FASEB J. 2019;33(4):5082–8.

75. Witcher KG, Eiferman DS, Godbout JP. Priming the Inflammatory Pump of the CNS after Traumatic Brain Injury. Trends Neurosci. 2015 Oct;38(10):609–20.

76. Papa L, Silvestri S, Brophy GM, Giordano P, Falk JL, Braga CF, et al. GFAP Out-Performs S100β in Detecting Traumatic Intracranial Lesions on Computed Tomography in Trauma Patients with Mild Traumatic Brain Injury and Those with Extracranial Lesions. J Neurotrauma. 2014 Nov 15;31(22):1815–22.

77. Papa L, Lewis LM, Falk JL, Zhang Z, Silvestri S, Giordano P, et al. Elevated levels of serum glial fibrillary acidic protein breakdown products in mild and moderate traumatic brain injury are associated with intracranial lesions and neurosurgical intervention. Ann Emerg Med. 2012 Jun;59(6):471–83.

78. Papa L, Brophy GM, Welch RD, Lewis LM, Braga CF, Tan CN, et al. Time Course and Diagnostic Accuracy of Glial and Neuronal Blood Biomarkers GFAP and UCH-L1 in a Large Cohort of Trauma Patients With and Without Mild Traumatic Brain Injury. JAMA Neurol. 2016 May 1;73(5):551–60.

79. Papa L, Zonfrillo MR, Welch RD, Lewis LM, Braga CF, Tan CN, et al. Evaluating glial and neuronal blood biomarkers GFAP and UCH-L1 as gradients of brain injury in concussive, subconcussive and non-concussive trauma: a prospective cohort study. BMJ Paediatr Open. 2019 Aug 1;3(1):e000473.

80. Žurek J, Fedora M. The usefulness of S100B, NSE, GFAP, NF-H, secretagogin and Hsp70 as a predictive biomarker of outcome in children with traumatic brain injury. Acta Neurochir (Wien). 2012 Jan 1;154(1):93–103.

81. Liddelow SA, Guttenplan KA, Clarke LE, Bennett FC, Bohlen CJ, Schirmer L, et al. Neurotoxic reactive astrocytes are induced by activated microglia. Nature. 2017 Jan;541(7638):481–7.

82. Xiong XY, Tang Y, Yang QW. Metabolic changes favor the activity and heterogeneity of reactive astrocytes. Trends Endocrinol Metab. 2022 Jun 1;33(6):390–400.

83. Gupte R, Brooks W, Vukas R, Pierce J, Harris J. Sex Differences in Traumatic Brain Injury: What We Know and What We Should Know. J Neurotrauma. 2019 Nov 15;36(22):3063–91.

84. Schäfer S, Wirths O, Multhaup G, Bayer TA. Gender dependent APP processing in a transgenic mouse model of Alzheimer’s disease. J Neural Transm. 2007 Mar 1;114(3):387– 94.

85. Jankowsky JL, Zheng H. Practical considerations for choosing a mouse model of Alzheimer’s disease. Mol Neurodegener. 2017 Dec 22;12(1):89.

86. Kapadia M, Mian MF, Michalski B, Azam AB, Ma D, Salwierz P, et al. Sex-Dependent Differences in Spontaneous Autoimmunity in Adult 3xTg-AD Mice. J Alzheimers Dis JAD. 2018;63(3):1191–205.

87. Prieto GA, Cotman CW. Cytokines and cytokine networks target neurons to modulate long-term potentiation. Cytokine Growth Factor Rev. 2017 Apr;34:27–33.

88. Talbot S, Foster SL, Woolf CJ. Neuroimmunity: Physiology and Pathology. Annu Rev Immunol. 2016;34(1):421–47.

89. Fukunaga K, Miyamoto E. Role of MAP kinase in neurons. Mol Neurobiol. 1998 Feb;16(1):79–95.

90. Asih PR, Prikas E, Stefanoska K, Tan ARP, Ahel HI, Ittner A. Functions of p38 MAP Kinases in the Central Nervous System. Front Mol Neurosci [Internet]. 2020 [cited 2023 May 3];13. Available from: https://www.frontiersin.org/articles/10.3389/fnmol.2020.570586

91. Schnöder L, Hao W, Qin Y, Liu S, Tomic I, Liu X, et al. Deficiency of Neuronal p38α MAPK Attenuates Amyloid Pathology in Alzheimer Disease Mouse and Cell Models through Facilitating Lysosomal Degradation of BACE1. J Biol Chem. 2016 Jan 29;291(5):2067–79.

92. Shih RH, Wang CY, Yang CM. NF-kappaB Signaling Pathways in Neurological Inflammation: A Mini Review. Front Mol Neurosci. 2015 Dec 18;8:77.

93. Di Cesare Mannelli L, Micheli L, Cervetto C, Toti A, Lucarini E, Parisio C, et al. Neuronal alarmin IL-1α evokes astrocyte-mediated protective signals: Effectiveness in chemotherapy-induced neuropathic pain. Neurobiol Dis. 2022 Jun 15;168:105716.

94. Michael BD, Bricio-Moreno L, Sorensen EW, Miyabe Y, Lian J, Solomon T, et al. Astrocyte- and Neuron-Derived CXCL1 Drives Neutrophil Transmigration and Blood-Brain Barrier Permeability in Viral Encephalitis. Cell Rep. 2020 Sep 15;32(11):108150.

95. Hallett H, Churchill L, Taishi P, De A, Krueger JM. Whisker stimulation increases expression of nerve growth factor- and interleukin-1beta-immunoreactivity in the rat somatosensory cortex. Brain Res. 2010 May 28;1333:48–56.

96. Ravizza T, Boer K, Redeker S, Spliet WGM, van Rijen PC, Troost D, et al. The IL-1β system in epilepsy-associated malformations of cortical development. Neurobiol Dis. 2006 Oct 1;24(1):128–43.

97. Meola D, Huang Z, Petitto JM. Selective Neuronal and Brain Regional Expession of IL-2 in IL2P 8-GFP Transgenic Mice: Relation to Sensorimotor Gating. J Alzheimers Dis Park. 2013 Oct 28;3(4):1000127.

98. Guo W, Zheng DH, Sun FJ, Yang JY, Zang ZL, Liu SY, et al. Expression and Cellular Distribution of the Interleukin 2 Signaling System in Cortical Lesions From Patients With Focal Cortical Dysplasia. J Neuropathol Exp Neurol. 2014 Mar 1;73(3):206–22.

99. Fontaine RH, Cases O, Lelièvre V, Mesplès B, Renauld JC, Loron G, et al. IL-9/IL-9 receptor signaling selectively protects cortical neurons against developmental apoptosis. Cell Death Differ. 2008 Oct;15(10):1542–52.

100. olde Heuvel F, Holl S, Chandrasekar A, Li Z, Wang Y, Rehman R, et al. STAT6 mediates the effect of ethanol on neuroinflammatory response in TBI. Brain Behav Immun. 2019 Oct 1;81:228–46.

101. Li S, olde Heuvel F, Rehman R, Aousji O, Froehlich A, Li Z, et al. Interleukin-13 and its receptor are synaptic proteins involved in plasticity and neuroprotection. Nat Commun. 2023 Jan 13;14(1):200.

102. Mori S, Sugama S, Nguyen W, Michel T, Sanna MG, Sanchez-Alavez M, et al. Lack of interleukin-13 receptor α1 delays the loss of dopaminergic neurons during chronic stress. J Neuroinflammation. 2017 Apr 21;14(1):88.

103. Xia M, Hyman BT. GROα/KC, a chemokine receptor CXCR2 ligand, can be a potent trigger for neuronal ERK1/2 and PI-3 kinase pathways and for tau hyperphosphorylation—a role in Alzheimer’s disease? J Neuroimmunol. 2002 Jan 1;122(1):55–64.

104. Carroll SL, Frohnert PW. Expression of JE (Monocyte Chemoattractant Protein-1) Is Induced by Sciatic Axotomy in Wild Type Rodents but Not in C57BL/Wlds Mice. J Neuropathol Exp Neurol. 1998 Oct 1;57(10):915–30.

105. Sankar SB, Pybus AF, Liew A, Sanders B, Shah KJ, Wood LB, et al. Low cerebral blood flow is a non-invasive biomarker of neuroinflammation after repetitive mild traumatic brain injury. Neurobiol Dis. 2019;124:544–54.

106. Lei J, Gao G, Feng J, Jin Y, Wang C, Mao Q, et al. Glial fibrillary acidic protein as a biomarker in severe traumatic brain injury patients: a prospective cohort study. Crit Care. 2015 Dec 1;19(1):362.

107. Thelin E, Al Nimer F, Frostell A, Zetterberg H, Blennow K, Nyström H, et al. A Serum Protein Biomarker Panel Improves Outcome Prediction in Human Traumatic Brain Injury. J Neurotrauma. 2019 Oct 15;36(20):2850–62.

108. McCrea M, Broglio SP, McAllister TW, Gill J, Giza CC, Huber DL, et al. Association of Blood Biomarkers With Acute Sport-Related Concussion in Collegiate Athletes: Findings From the NCAA and Department of Defense CARE Consortium. JAMA Netw Open. 2020 Jan 3;3(1):e1919771.

109. Xu X, Cowan M, Beraldo F, Schranz A, McCunn P, Geremia N, et al. Repetitive mild traumatic brain injury in mice triggers a slowly developing cascade of long-term and persistent behavioral deficits and pathological changes. Acta Neuropathol Commun. 2021 Apr 6;9(1):60.

110. Button EB, Cheng WH, Barron C, Cheung H, Bashir A, Cooper J, et al. Development of a novel, sensitive translational immunoassay to detect plasma glial fibrillary acidic protein (GFAP) after murine traumatic brain injury. Alzheimers Res Ther. 2021 Mar 7;13(1):58.

111. Shandra O, Winemiller AR, Heithoff BP, Munoz-Ballester C, George KK, Benko MJ, et al. Repetitive Diffuse Mild Traumatic Brain Injury Causes an Atypical Astrocyte Response and Spontaneous Recurrent Seizures. J Neurosci Off J Soc Neurosci. 2019 Mar 6;39(10):1944–63.

112. Singh K, Trivedi R, Devi MM, Tripathi RP, Khushu S. Longitudinal changes in the DTI measures, anti-GFAP expression and levels of serum inflammatory cytokines following mild traumatic brain injury. Exp Neurol. 2016 Jan 1;275:427–35.

113. Michinaga S, Koyama Y. Pathophysiological Responses and Roles of Astrocytes in Traumatic Brain Injury. Int J Mol Sci. 2021 Jun 15;22(12):6418.

114. Villapol S, Byrnes KR, Symes AJ. Temporal dynamics of cerebral blood flow, cortical damage, apoptosis, astrocyte-vasculature interaction and astrogliosis in the pericontusional region after traumatic brain injury. Front Neurol. 2014;5:82.

115. Todd BP, Chimenti MS, Luo Z, Ferguson PJ, Bassuk AG, Newell EA. Traumatic brain injury results in unique microglial and astrocyte transcriptomes enriched for type I interferon response. J Neuroinflammation. 2021 Jul 5;18:151.

116. Arneson D, Zhang G, Ying Z, Zhuang Y, Byun HR, Ahn IS, et al. Single cell molecular alterations reveal target cells and pathways of concussive brain injury. Nat Commun. 2018 Sep 25;9(1):3894.

117. Choudhury GR, Ryou MG, Poteet E, Wen Y, He R, Sun F, et al. INVOLVEMENT OF P38 MAPK IN REACTIVE ASTROGLIOSIS INDUCED BY ISCHEMIC STROKE. Brain Res. 2014 Mar 3;1551:45.

118. Mandell JW, VandenBerg SR. ERK/MAP kinase is chronically activated in human reactive astrocytes. Neuroreport. 1999 Nov 26;10(17):3567–72.

119. Kaminska B, Gozdz A, Zawadzka M, Ellert-Miklaszewska A, Lipko M. MAPK Signal Transduction Underlying Brain Inflammation and Gliosis as Therapeutic Target. Anat Rec. 2009;292(12):1902–13.

120. Oddo S, Caccamo A, Shepherd JD, Murphy MP, Golde TE, Kayed R, et al. Triple-transgenic model of Alzheimer’s disease with plaques and tangles: intracellular Abeta and synaptic dysfunction. Neuron. 2003 Jul 31;39(3):409–21.

121. Billings LM, Oddo S, Green KN, McGaugh JL, LaFerla FM. Intraneuronal Abeta causes the onset of early Alzheimer’s disease-related cognitive deficits in transgenic mice. Neuron. 2005 Mar 3;45(5):675–88.

