## Supplementary Information for "Profiling the neuroimmune cascade in 3xTg mice exposed to successive mild traumatic brain injuries"

| | |
|---|---|
| **Erin M. Buckley**, PhD[¥,] *<br>Assistant Professor<br>Wallace H. Coulter Department of Biomedical Engineering, Georgia Institute of Technology and Emory University; Department of Pediatrics, School of Medicine, Emory University<br>1760 Haygood Dr NE<br>Atlanta, GA 30322<br> | **Levi B. Wood**, PhD [¥,] *<br>Assistant Professor<br>George W. Woodruff School of Mechanical Engineering, Wallace H. Coulter Department of Biomedical Engineering, and Parker H. Petit Institute for Bioengineering and Bioscience, Georgia Institute of Technology<br>315 Ferst Dr, Rm 3311<br>Atlanta, GA 30332<br> |

**Supplementary Table S1. Sample sizes of each experimental group.** Tables are separated by Female (top) and Male (bottom) samples by injury group (1x,3x,5xCHI) and time point (pre-injury, 30min, 4hr, 24hr). Sample size is indicated in n column. In total, 96 females and 83 males were used in the protein analysis and 39 females in RNAseq analysis.

### Protein Data

| Sex | Injury | Time Point | Total Time | n |
|---|---|---|---|---|
| Females | 1xCHI | Pre-Injury | 0 | 10 |
| | | 30min | 0.5 | 8 |
| | | 4hr | 4 | 8 |
| | | 24hr | 24 | 8 |
| | 3xCHI | Pre-Injury | 48 | 8 |
| | | 30min | 48.5 | 8 |
| | | 4hr | 52 | 6 |
| | | 24hr | 72 | 8 |
| | 5xCHI | Pre-Injury | 96 | 8 |
| | | 30min | 96.5 | 8 |
| | | 4hr | 100 | 7 |
| | | 24hr | 120 | 9 |
| | | | Female Total: | **96** |
| Males | 1xCHI | Pre-Injury | 0 | 13 |
| | | 30min | 0.5 | 7 |
| | | 4hr | 4 | 6 |
| | | 24hr | 24 | 6 |
| | 3xCHI | Pre-Injury | 48 | 7 |
| | | 30min | 48.5 | 6 |
| | | 4hr | 52 | 7 |
| | | 24hr | 72 | 7 |
| | 5xCHI | Pre-Injury | 96 | 6 |
| | | 30min | 96.5 | 6 |
| | | 4hr | 100 | 6 |
| | | 24hr | 120 | 6 |
| | | | Male Total: | **83** |

### RNAseq Data

| Sex | Injury | Time Point | Total Time | n |
|---|---|---|---|---|
| Females | 1xCHI | Pre-Injury | 0 | 4 |
| | | 30min | 0.5 | 3 |
| | | 4hr | 4 | 3 |
| | | 24hr | 24 | 4 |
| | 3xCHI | Pre-Injury | 48 | 3 |
| | | 30min | 48.5 | 3 |
| | | 4hr | 52 | 2 |
| | | 24hr | 72 | 4 |
| | 5xCHI | Pre-Injury | 96 | 3 |
| | | 30min | 96.5 | 3 |
| | | 4hr | 100 | 3 |
| | | 24hr | 120 | 4 |
| | | | Female Total: | **39** |

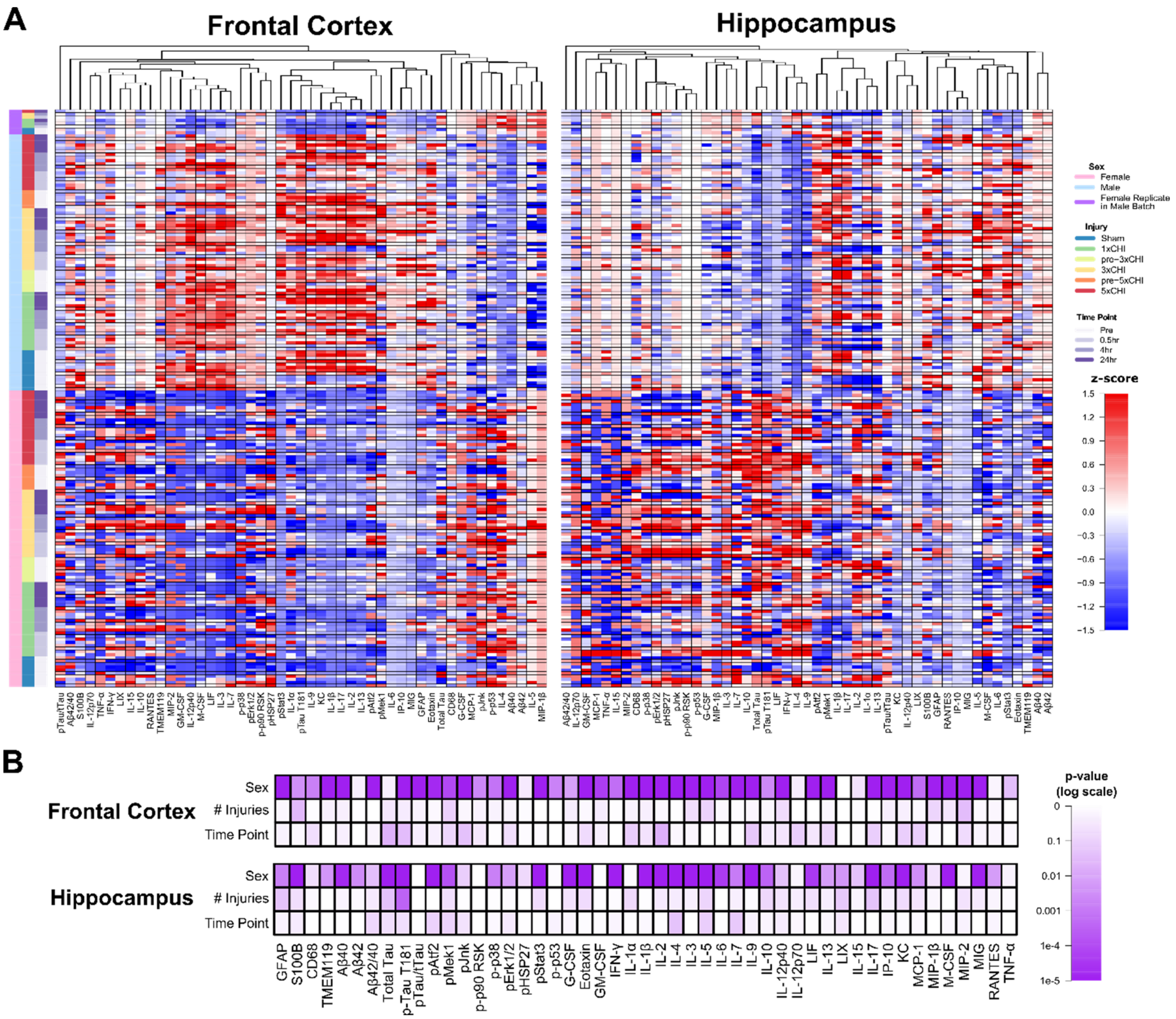


**Supplementary Figure S1. Pronounced sexual dimorphism of immune signaling, glial markers, and molecular markers of pathology in 3xTg-AD mice: all-samples heatmap and multiple linear regression model. A**) 30 cytokines, 6 MAPK phospho-proteins, amyloid beta 40 and 42, total and phospho-tau, GFAP, S100B, CD68, and TMEM119 were measured by multiplex and traditional ELISA in both frontal cortex (left) and hippocampus (right) tissue of 96 female (pink) and 83 male (blue) 3xTg-AD mice aged 2-4mo. Eight female samples were run as replicates within male batch assays (purple) to allow transformation of male data to female space by linear regression. S100B, CD68, and TMEM119 were run with all samples at the same time while all other assays were run in separate batches for male and female samples. Each row represents data from an individual mouse z-scored along each column. Columns are clustered by Euclidean distance via unweighted pair group method with arithmetic mean. **B**) Multiple linear regression models were calculated for each measured protein using fixed effects of sex, number of injuries, and time point after most recent injury. The model shows that the effect of sex strongly outweighs injury-dependent changes for most proteins from both regions.

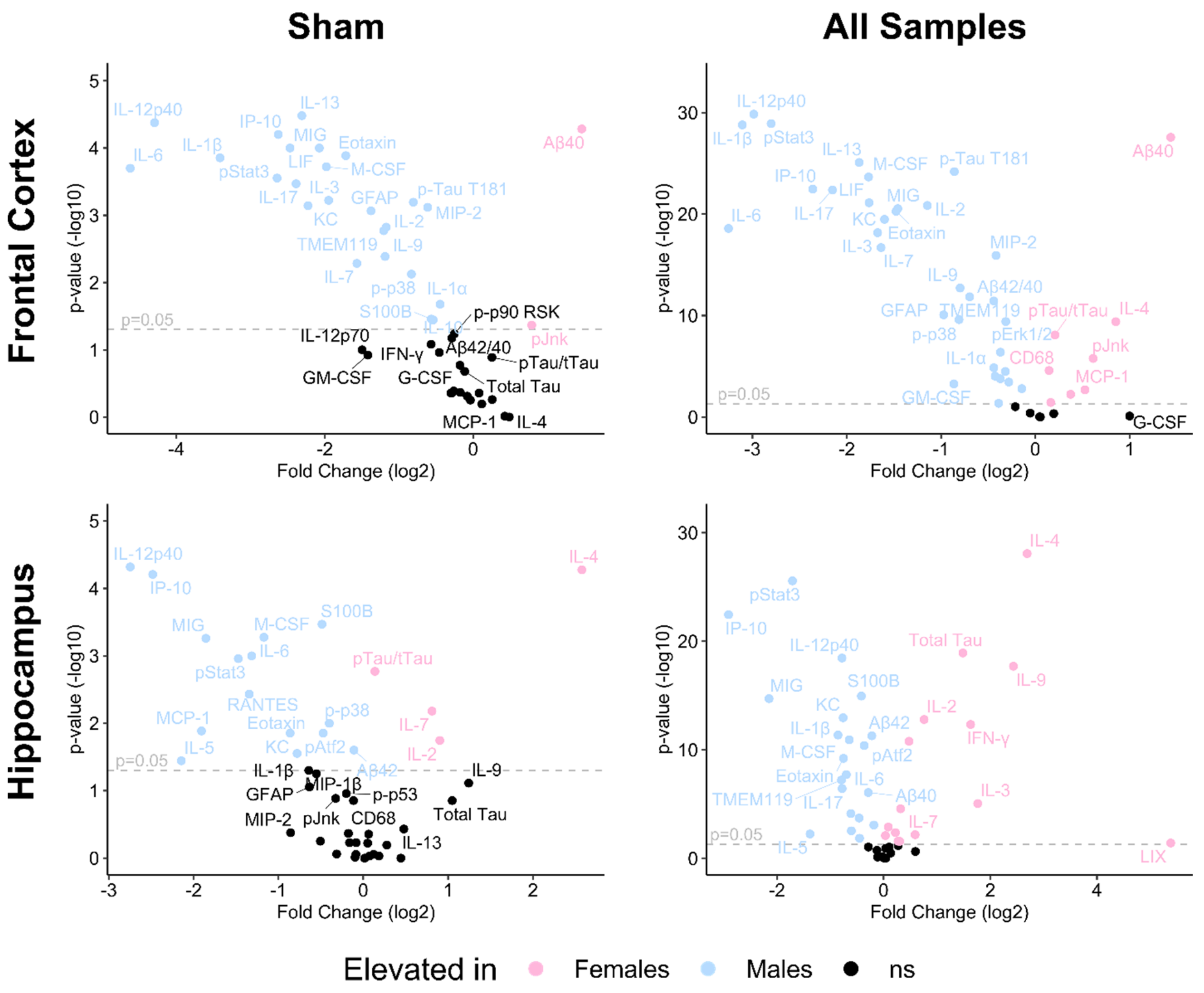


**Supplementary Figure S2. Pronounced sexual dimorphism of immune signaling, glial markers, and molecular markers of pathology in 3xTg-AD mice: volcano plots.** Volcano plots of male vs female comparative analysis show Wilcoxon rank-sum p-values (vertical axis, -log10 scale) and fold change (horizontal axis, log2 scale) of each measured protein. Proteins are considered significantly elevated in either male (blue) or female (pink) samples if $p<0.05$ by Wilcoxon rank-sum test. All significantly different proteins within sham-injured samples were also significant within all samples in the same direction. The all-samples analysis revealed a greater proportion of significantly different proteins by sex.

**Supplementary Table S2. Sex differences by protein within each brain region among sham-injured and all samples.** Proteins are considered significantly elevated in either male (blue) or female (pink) samples if p<0.05 by Wilcoxon rank-sum test. All significantly different proteins within sham-injured samples were also significant within all samples in the same direction. ns = not significant, *p<0.05, **p<0.01, ***p<0.001, ****p<0.0001. The all-samples analysis revealed a greater proportion of significantly different proteins by sex.

| Protein | Frontal Cortex Sham | Frontal Cortex All | Hippocampus Sham | Hippocampus All |
|---|---|---|---|---|
| GFAP | Males *** | Males **** | ns | Males * |
| S100B | Males * | Males *** | Males *** | Males **** |
| CD68 | ns | Females **** | ns | ns |
| TMEM119 | Males ** | Males **** | ns | Males **** |
| Aβ40 | Females **** | Females **** | ns | Males **** |
| Aβ42 | ns | Females * | Males * | Males **** |
| Total Tau | ns | ns | ns | Females **** |
| p-Tau | Males *** | Males **** | ns | Females **** |
| pAtf2 | ns | Males **** | Males * | Males **** |
| pMek1 | ns | Males **** | ns | Males *** |
| pJnk | Females * | Females **** | ns | ns |
| p-p90 RSK | ns | Males ** | N/A | N/A |
| p-p38 | Males ** | Males **** | Males * | Males *** |
| pErk1/2 | ns | Males **** | ns | Females ** |
| pHSP27 | ns | ns | ns | ns |
| pStat3 | Males *** | Males **** | Males ** | Males **** |
| p-p53 | ns | Females ** | ns | ns |
| G-CSF | ns | ns | Females ** | Females **** |
| Eotaxin | Males *** | Males **** | Males * | Males **** |
| GM-CSF | ns | Males *** | ns | Females ** |
| IFN-γ | ns | Males *** | ns | Females **** |
| IL-1α | Males * | Males **** | ns | ns |
| IL-1β | Males *** | Males **** | ns | Males **** |
| IL-2 | Males ** | Males **** | Females * | Females **** |
| IL-4 | ns | Females **** | Females **** | Females **** |
| IL-3 | Males *** | Males **** | Females * | Females **** |
| IL-5 | Females * | Females **** | Males * | Males ** |
| IL-6 | Males *** | Males **** | Males ** | Males **** |
| IL-7 | Males ** | Males **** | Females ** | Females **** |
| IL-9 | Males ** | Males **** | ns | Females **** |
| IL-10 | Males * | Males **** | ns | Females ** |
| IL-12p40 | Males **** | Males **** | Males **** | Males **** |
| IL-12p70 | ns | Ns | ns | ns |
| LIF | Males *** | Males **** | ns | ns |
| IL-13 | Males **** | Males **** | ns | Females * |
| LIX | N/A | N/A | ns | Females * |
| IL-15 | ns | ns | ns | ns |
| IL-17 | Males *** | Males **** | ns | Males **** |
| IP-10 | Males **** | Males **** | Males **** | Males **** |
| KC | Males *** | Males **** | Males * | Males **** |
| MCP-1 | ns | Females ** | Males * | Males **** |
| MIP-1β | Females ** | Females **** | ns | Females * |
| M-CSF | Males *** | Males **** | Males *** | Males **** |
| MIP-2 | Males *** | Males **** | ns | Females ** |
| MIG | Males *** | Males **** | Males *** | Males **** |
| RANTES | ns | ns | Males ** | Males ** |
| TNF-α | ns | Males * | ns | ns |

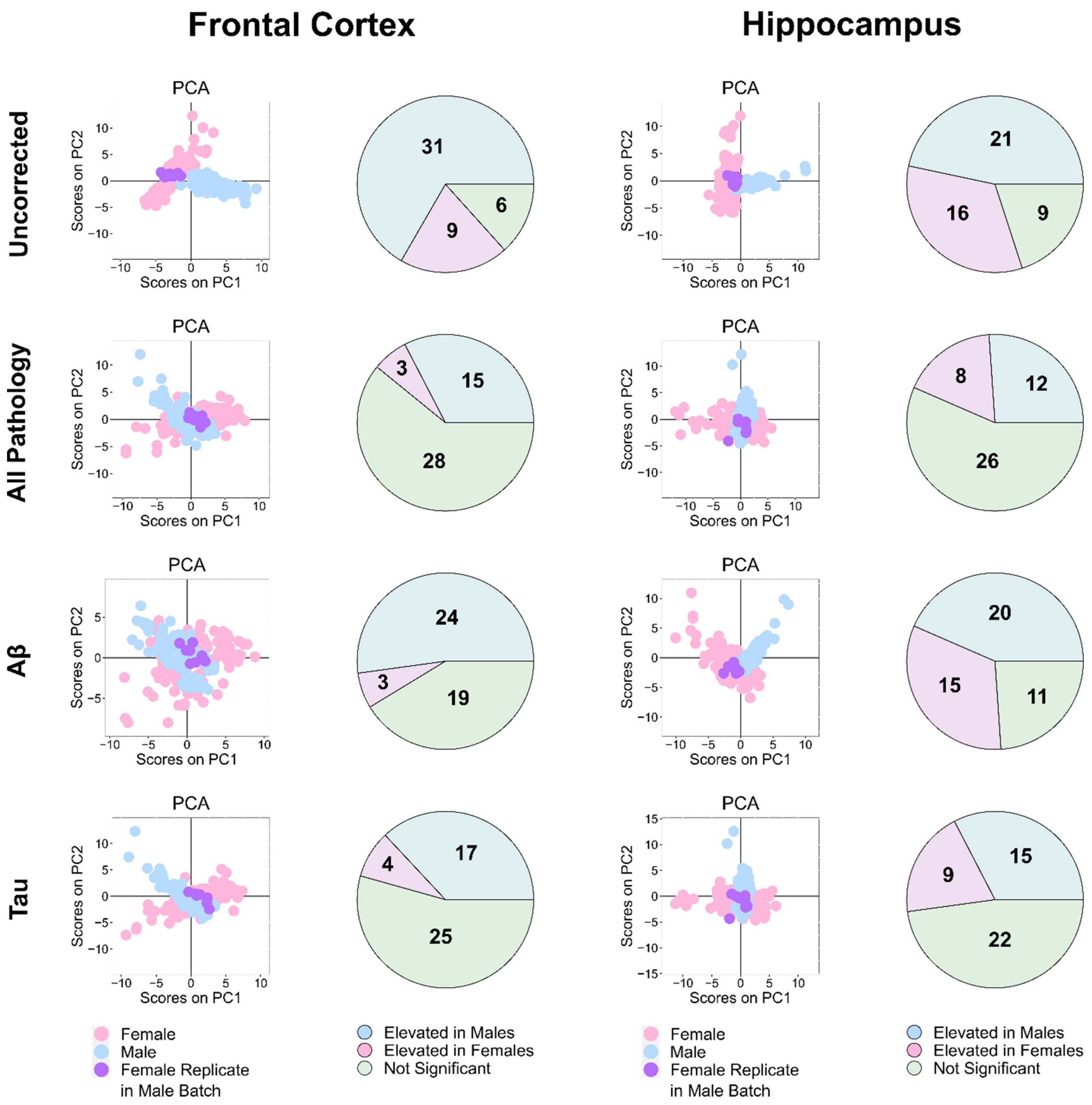


**Supplementary Figure S3. Sex differences within pathology-adjusted data.** Adjusted data is defined as the residual of a linear regression of Aβ markers (Aβ40 + Aβ42), tau markers (total tau + phospho-tau T181), or all pathological markers (Aβ and tau combined) against all combined protein data. From top to bottom, the graphic shows uncorrected data, all pathology-, Aβ-, and tau-adjusted data for both the frontal cortex (left) and hippocampus (right). A principal component analysis (PCA) scores plot of the first two PCs is shown to illustrate changes in variance between the samples after adjustment. The pie charts are annotated with the number of differentially expressed proteins which are significantly upregulated in male (blue) and female (pink) samples, or not significantly different (green). Proteins are considered significantly different if $p<0.05$ by Wilcox Rank Sum test.

**Supplementary Table S3. Sex differences by protein within each brain region before and after adjustment for pathological markers.** Proteins are considered significantly elevated in either male (blue) or female (pink) samples if $p<0.05$ by Wilcoxon rank-sum test. Adjusted data is defined as the residual of a linear regression of Aβ markers (Aβ40 + Aβ42), tau markers (total tau + phospho-tau T181), or all pathological markers (Aβ and tau combined) against all combined protein data. Phospho-p90 RSK and LIX were measured below the detection limit among all eight female replicate samples within the male batch hippocampus and frontal cortex, respectively, disallowing conversion to female space.

| | Frontal Cortex | | | | Hippocampus | | | |
|---|---|---|---|---|---|---|---|---|
| Protein | Original | All | Aβ | Tau | Original | All | Aβ | Tau |
| GFAP | M | ns | ns | M | M | ns | ns | ns |
| S100B | M | ns | ns | ns | M | M | M | M |
| CD68 | F | F | F | F | ns | ns | ns | ns |
| TMEM119 | M | M | ns | M | M | ns | M | ns |
| Aβ40 | F | F | M | F | M | M | M | M |
| Aβ42 | F | M | M | M | M | F | ns | M |
| Total Tau | ns | ns | ns | M | F | ns | F | M |
| Phospho-Tau | M | M | M | ns | F | ns | F | M |
| pAtf2 | M | ns | M | ns | M | M | M | M |
| pMek1 | M | ns | ns | ns | M | M | M | M |
| pJnk | F | ns | ns | ns | ns | ns | ns | ns |
| p-p90 RSK | M | ns | ns | ns | N/A | N/A | N/A | N/A |
| p-p38 | M | M | M | M | M | ns | M | M |
| pErk1/2 | M | M | M | M | F | ns | F | ns |
| pHSP27 | ns | ns | ns | ns | ns | ns | F | ns |
| pStat3 | M | ns | M | ns | M | M | M | M |
| p-p53 | F | ns | ns | ns | ns | ns | ns | ns |
| G-CSF | ns | ns | ns | ns | F | ns | F | ns |
| Eotaxin | M | ns | ns | ns | M | ns | M | ns |
| GM-CSF | M | M | M | M | F | M | M | F |
| IFN-γ | M | ns | M | ns | F | F | F | F |
| IL-1α | M | ns | ns | ns | ns | ns | ns | ns |
| IL-1β | M | ns | M | ns | M | M | M | M |
| IL-2 | M | ns | M | ns | F | F | F | F |
| IL-4 | F | ns | F | F | F | F | F | F |
| IL-3 | M | M | M | M | F | F | F | F |
| IL-5 | F | ns | ns | ns | M | M | M | M |
| IL-6 | M | ns | ns | M | M | ns | M | ns |
| IL-7 | M | M | M | M | F | ns | F | ns |
| IL-9 | M | ns | M | ns | F | F | F | F |
| IL-10 | M | M | M | M | F | ns | F | ns |
| IL-12p40 | M | M | M | M | M | ns | M | ns |
| IL-12p70 | ns | M | ns | M | ns | ns | ns | ns |
| LIF | M | M | M | M | ns | F | ns | F |
| IL-13 | M | ns | M | ns | F | ns | F | ns |
| LIX | N/A | N/A | N/A | N/A | F | ns | F | ns |
| IL-15 | ns | ns | ns | ns | ns | ns | ns | ns |
| IL-17 | M | ns | M | ns | M | M | M | M |
| IP-10 | M | M | ns | ns | M | ns | M | ns |
| KC | M | ns | M | ns | M | M | M | M |
| MCP-1 | F | ns | ns | ns | M | M | M | M |
| MIP-1β | F | F | F | F | F | ns | ns | F |
| M-CSF | M | M | M | M | M | M | M | M |
| MIP-2 | M | ns | M | M | F | F | F | F |
| MIG | M | ns | ns | M | M | ns | M | ns |
| RANTES | ns | M | M | ns | M | ns | M | ns |
| TNF-α | M | ns | M | ns | ns | ns | ns | ns |

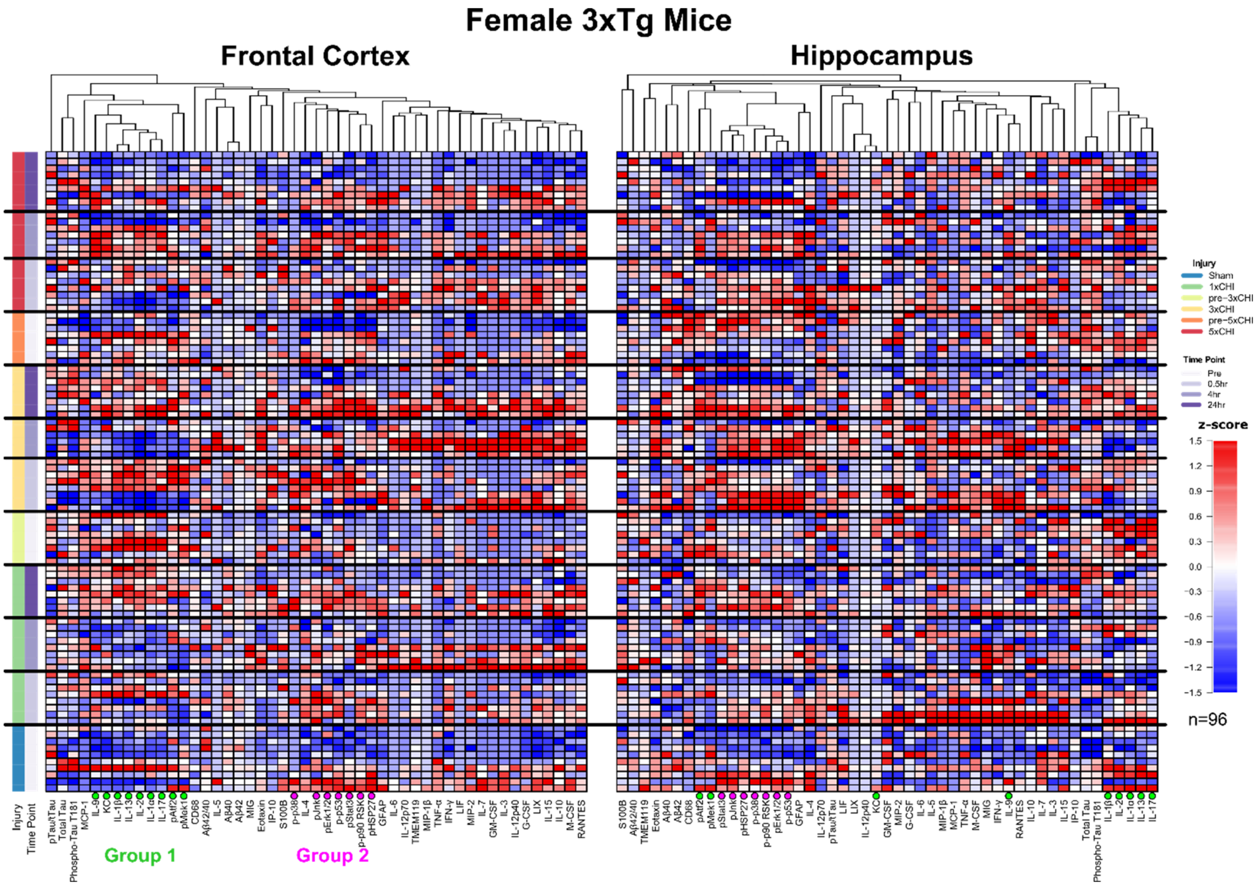


**Supplementary Figure S4. Female protein data: Quantification of immune and pathological markers in female 3xTg-AD mice pre- and 30min, 4hr, & 24hr post-1x, 3x, and 5xCHI.** 30 cytokines, 6 MAPK phospho-proteins, amyloid beta 40 and 42, total and phospho-tau, GFAP, S100B, CD68, and TMEM119 were measured by multiplex and traditional ELISA in both frontal cortex (left) and hippocampus (right) tissue of 96 female 3xTg-AD mice aged 2-4mo. Each row represents data from an individual mouse z-scored along each column. Columns are clustered by Euclidean distance via unweighted pair group method with arithmetic mean.

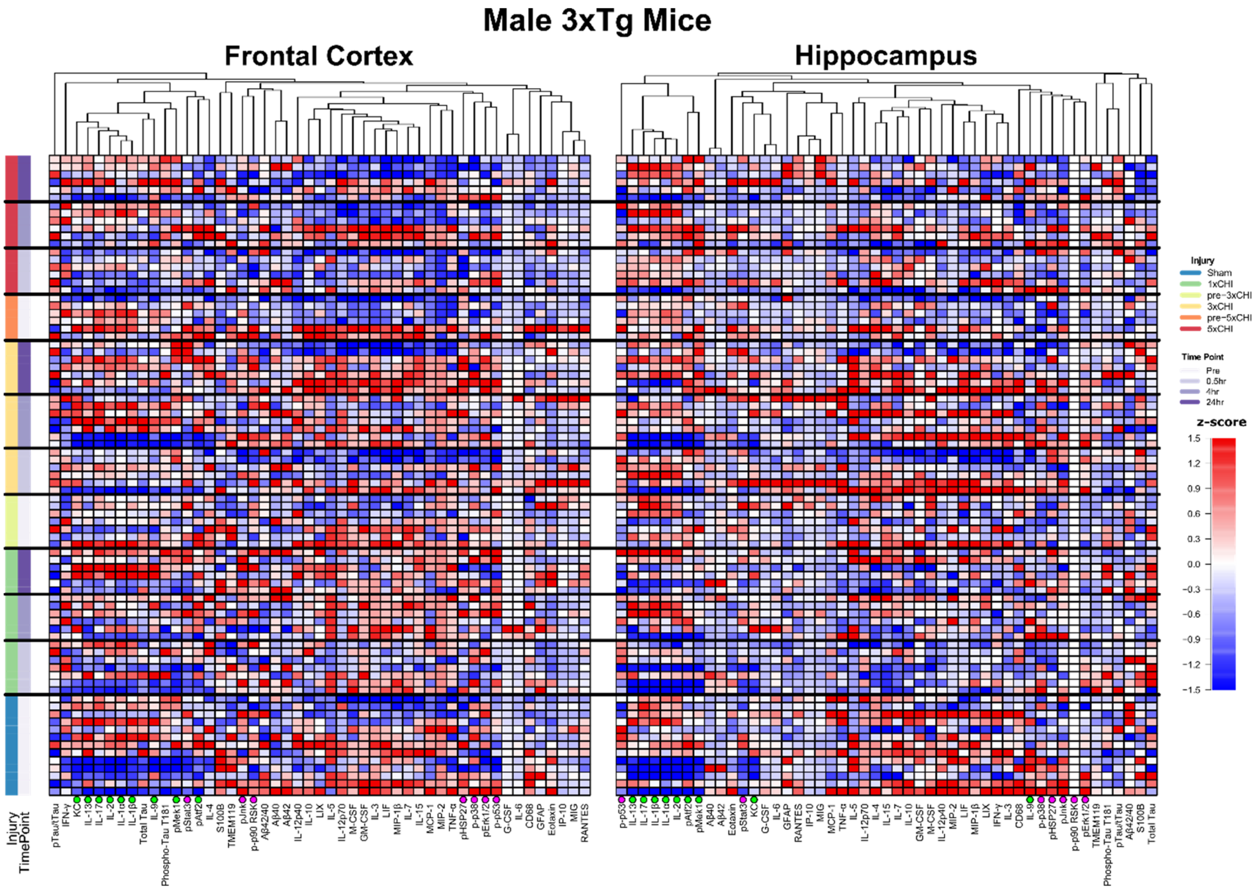


**Supplementary Figure S5. Male protein data: Quantification of immune and pathological markers in male 3xTg-AD mice pre- and 30min, 4hr, & 24hr post-1x, 3x, and 5xCHI.** 30 cytokines, 6 MAPK phospho-proteins, amyloid beta 40 and 42, total and phospho-tau, GFAP, S100B, CD68, and TMEM119 were measured by multiplex and traditional ELISA in both frontal cortex (left) and hippocampus (right) tissue of 83 male 3xTg-AD mice aged 2-4mo. Each row represents data from an individual mouse z-scored along each column. Columns are clustered by Euclidean distance via unweighted pair group method with arithmetic mean.

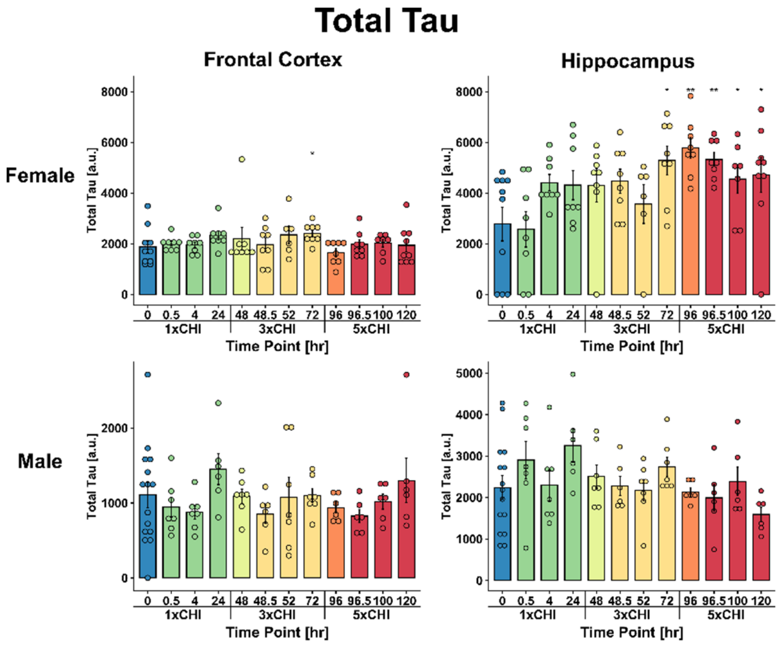


**Supplementary Figure S6. Total tau by sex and brain region.** Total tau was measured by Luminex ELISA (Millipore Sigma HNABTMAG-68K) in male and female frontal cortex and hippocampus tissue (*p<0.05, **p<0.01, Wilcox test vs sham) (mean ± SEM). Male and female samples run separately.

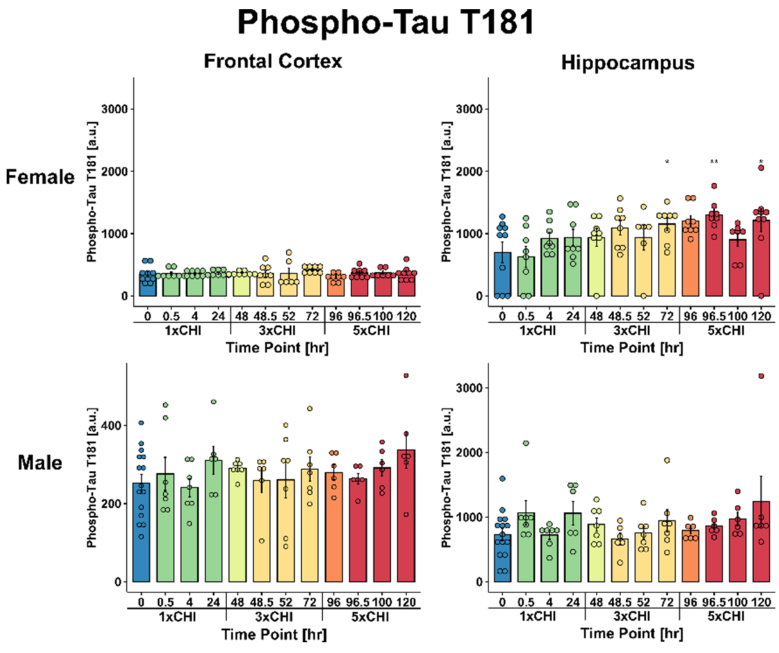


**Supplementary Figure S7. Phospho-tau T181 by sex and brain region.** Phospho-tau T181 was measured by Luminex ELISA (Millipore Sigma HNABTMAG-68K) in male and female frontal cortex and hippocampus tissue (*p<0.05, **p<0.01, Wilcox test vs sham) (mean ± SEM). Male and female samples run separately.

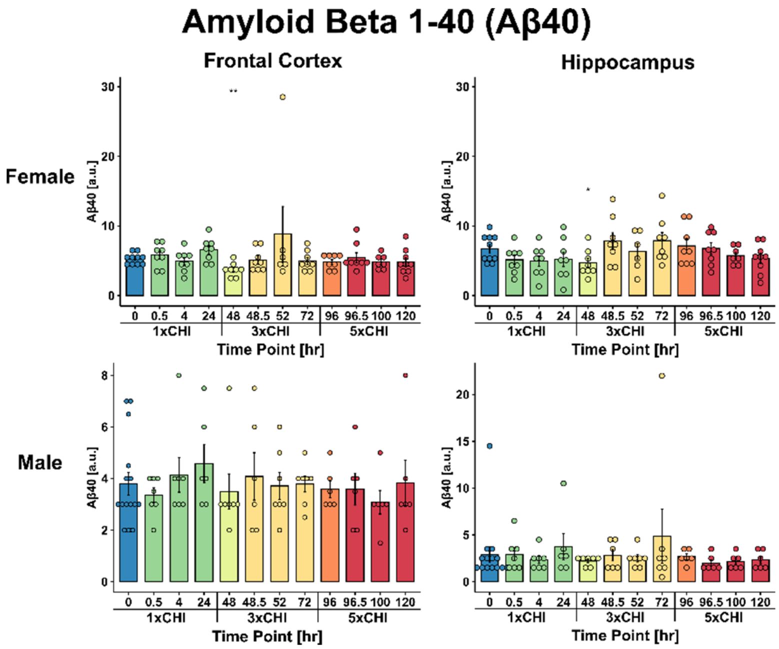


**Supplementary Figure S8. Amyloid Beta 1-40 (Aβ40) by sex and brain region.** Aβ40 was measured by Luminex ELISA (Millipore Sigma HNABTMAG-68K) in male and female frontal cortex and hippocampus tissue (*p<0.05, **p<0.01, Wilcox test vs sham) (mean ± SEM). Male and female samples run separately.

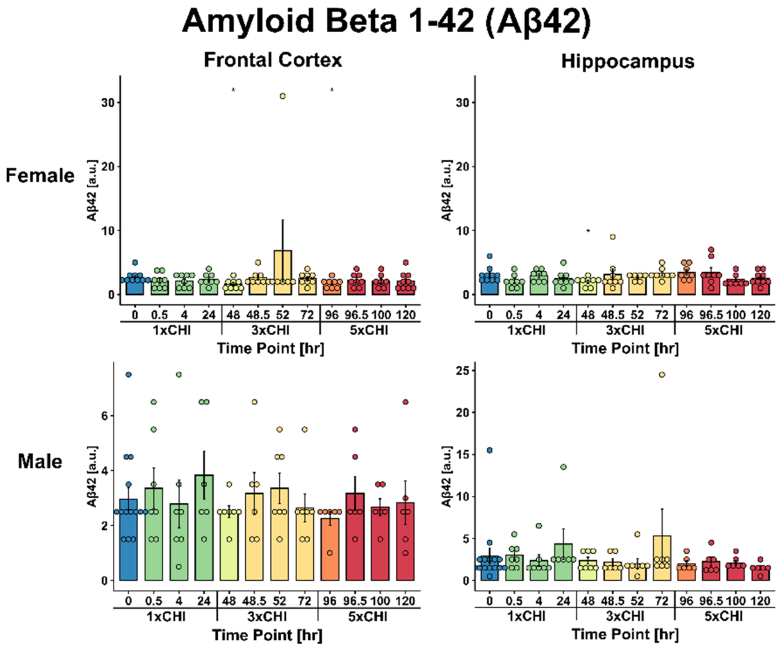


**Supplementary Figure S9. Amyloid Beta 1-42 (Aβ42) by sex and brain region.** Aβ42 was measured by Luminex ELISA (Millipore Sigma HNABTMAG-68K) in male and female frontal cortex and hippocampus tissue (*p<0.05, Wilcox test vs sham) (mean ± SEM). Male and female samples run separately.

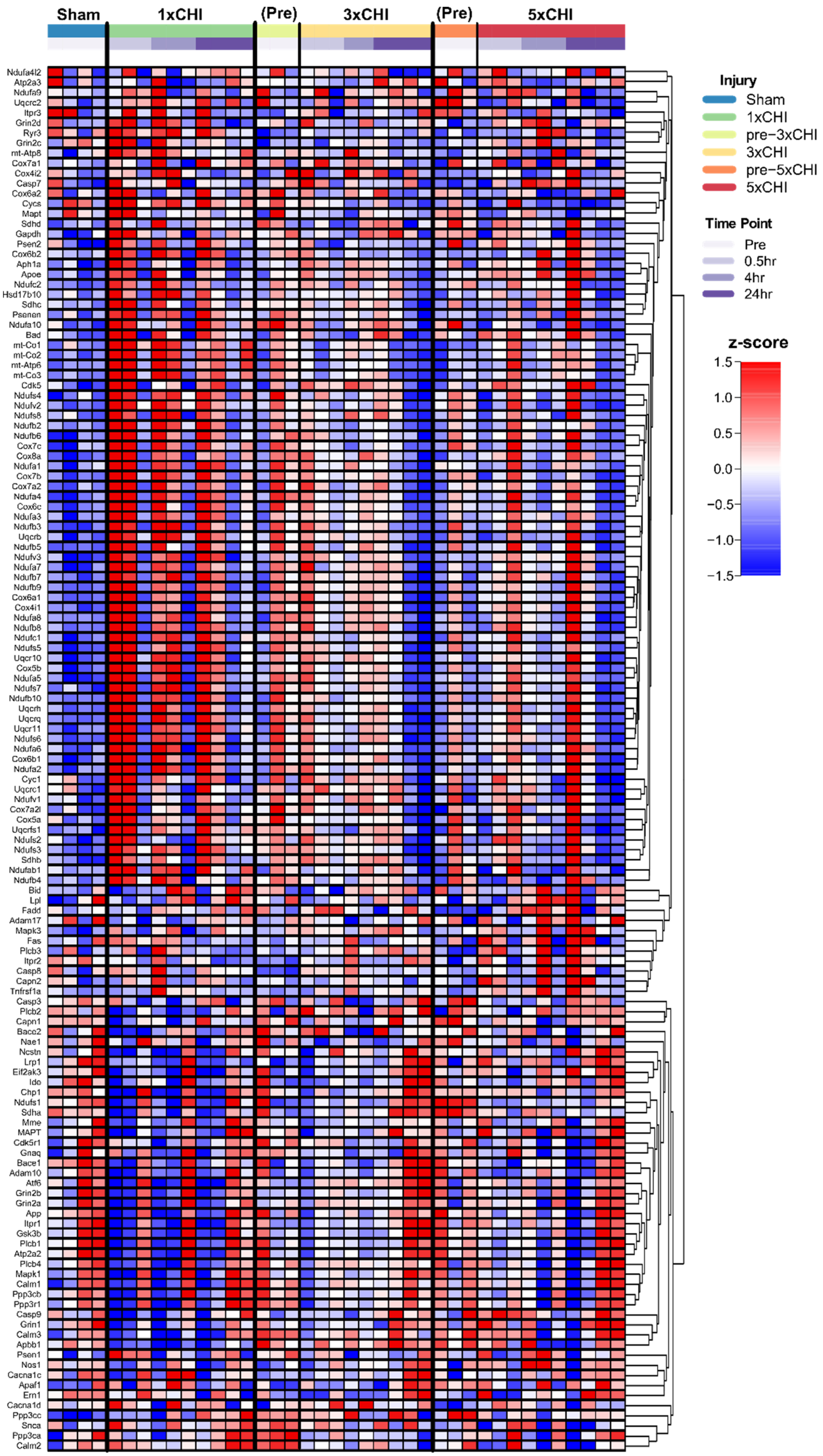

Sham
1xCHI
(Pre)
3xCHI
(Pre)
5xCHI
Ndufa4l2
Atp2a3
Ndufa9
Uqcrc2
Itpr3
Grin2d
Ryr3
Grin2c
mt-Atp8
Cox7a1
Cox4i2
Casp7
Cox6a2
Cycs
Mapt
Sdhd
Gapdh
Psen2
Cox6b2
Aph1a
Apoe
Ndufc2
Hsd17b10
Sdhc
Psenen
Ndufa10
Bad
mt-Co1
mt-Co2
mt-Atp6
mt-Co3
Cdk5
Ndufs4
Ndufv2
Ndufs8
Ndufb2
Ndufb6
Cox7c
Cox8a
Ndufa1
Cox7b
Cox7a2
Ndufa4
Cox6c
Ndufa3
Ndufb3
Uqcrb
Ndufb5
Ndufv3
Ndufa7
Ndufb7
Ndufb9
Cox6a1
Cox4i1
Ndufa8
Ndufb8
Ndufc1
Ndufs5
Uqcr10
Cox5b
Ndufa5
Ndufs7
Ndufb10
Uqcrh
Uqcrq
Uqcr11
Ndufs6
Ndufa6
Cox6b1
Ndufa2
Cyc1
Uqcrc1
Ndufv1
Cox7a2l
Cox5a
Uqcrfs1
Ndufs2
Ndufs3
Sdhb
Ndufab1
Ndufb4
Bid
Lpl
Fadd
Adam17
Mapk3
Fas
Plcb3
Itpr2
Casp8
Capn2
Tnfrsf1a
Casp3
Plcb2
Capn1
Bace2
Nae1
Ncstn
Lrp1
Eif2ak3
Ide
Chp1
Ndufs1
Sdha
Mme
MAPT
Cdk5r1
Gnaq
Bace1
Adam10
Atf6
Grin2b
Grin2a
App
Itpr1
Gsk3b
Plcb1
Atp2a2
Plcb4
Mapk1
Calm1
Ppp3cb
Ppp3r1
Casp9
Grin1
Calm3
Apbb1
Psen1
Nos1
Cacna1c
Apaf1
Ern1
Cacna1d
Ppp3cc
Snca
Ppp3ca
Calm2
Injury
Sham
1xCHI
pre−3xCHI
3xCHI
pre−5xCHI
5xCHI
Time Point
Pre
0.5hr
4hr
24hr
z-score
1.5
1.0
0.5
0.0
−0.5
−1.0
−1.5

**Supplementary Figure S10. Kyoto Encyclopedia of Genes and Genomes (KEGG) Alzheimer's disease gene set.** RNAseq data from 204mo female 3xTg somato-motor cortex lysate pre- and 30min, 4hr, and 24hr post- 1x, 3x, and 5xCHI (n=2-4). Heatmap shows subset of all expressed genes from the KEGG Alzheimer's disease gene set (MSigDB C2 collection). RNAseq data is normalized, variance stabilized (**Methods**), then z-scored within each gene (rows) across all animals (columns). Genes are clustered by unweighted pair group method with arithmetic mean using Euclidean distance.

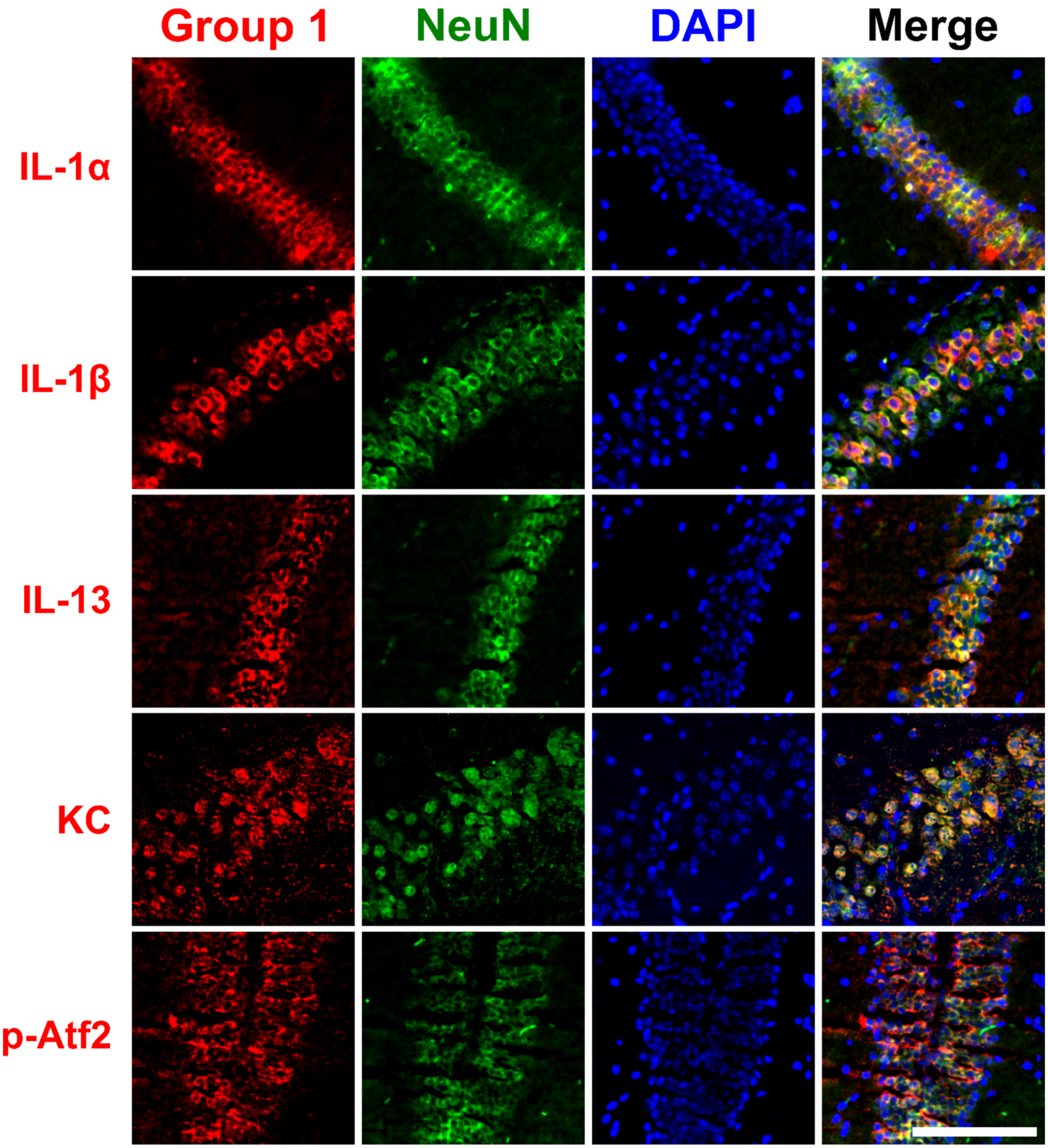


**Supplementary Figure S11. Group 1 cytokines and phospho-proteins co-localize with NeuN in hippocampus 24hr post-CHI.** Group 1 cytokines (IL-1α, IL-1β, IL-13, and KC) and MAPK phospho-protein phospho-Atf2 (red) co-stained with NeuN (green) and DAPI (blue) show extensive neuronal localization in the hippocampus 24h post-CHI in female 3xTg mice aged 2-4mo (scale bar: 500μm).

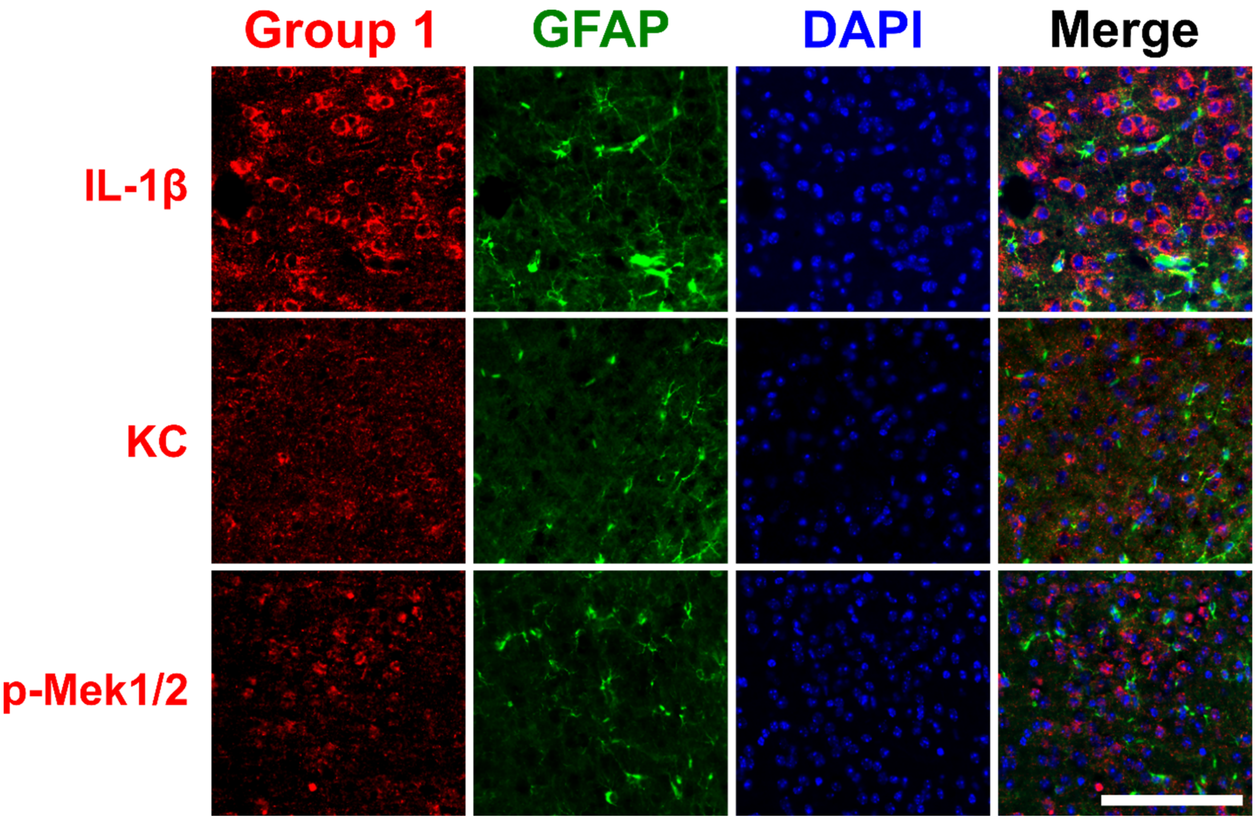


**Supplementary Figure S12. Group 1 cytokines and phospho-proteins do not co-localize with astrocyte marker GFAP in the cortex 24hr post-CHI.** Group 1 cytokines (IL-1β and KC) and MAPK phospho-protein phospho-Mek1/2 (red) co-stained with GFAP (green) and DAPI (blue) showed minimal astrocyte co-localization in the cortex 24h post-CHI in female 3xTg mice aged 2-4mo (scale bar: 500μm).

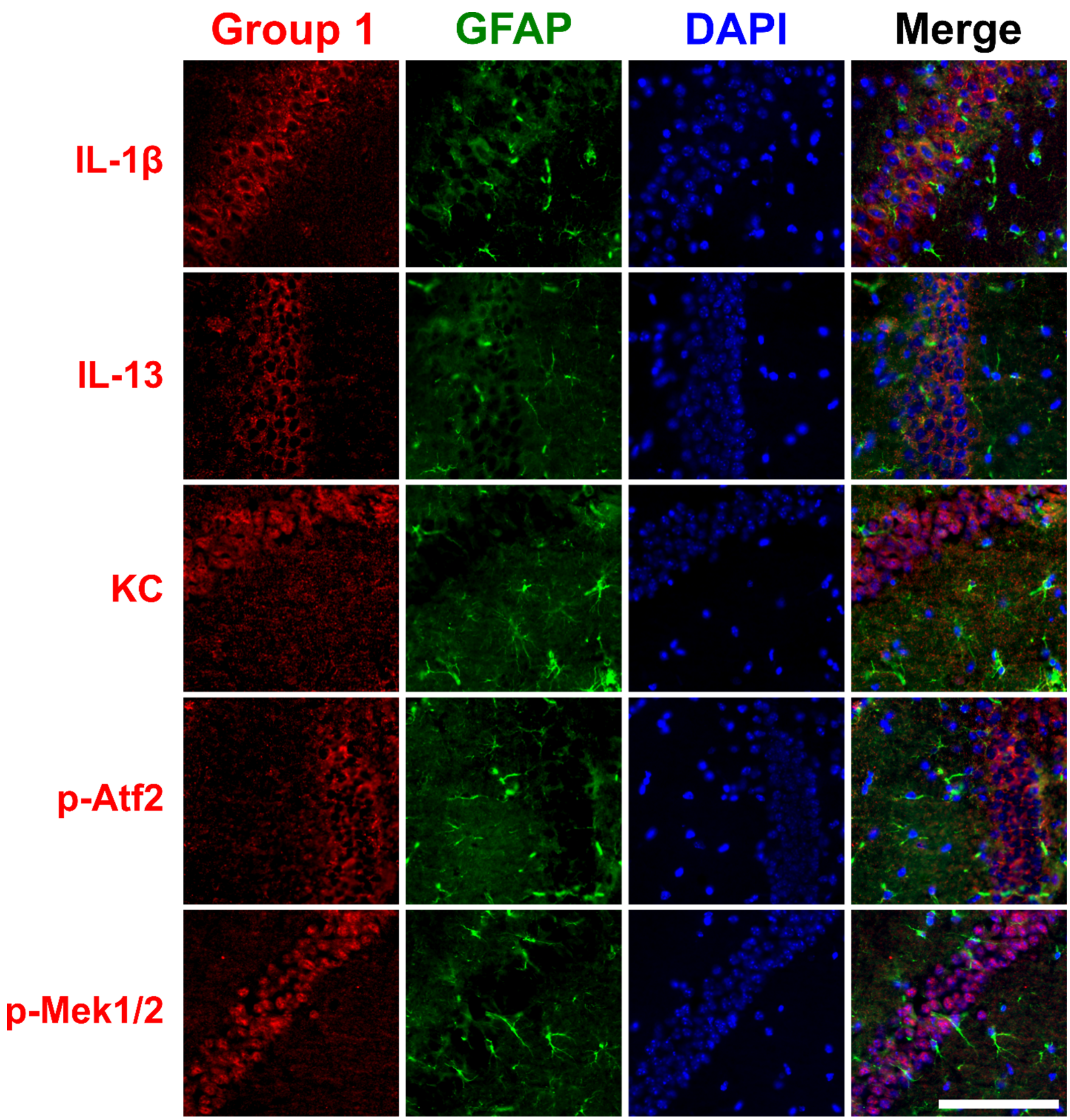


**Supplementary Figure S13. Group 1 cytokines and phospho-proteins do not co-localize with astrocyte marker GFAP in the hippocampus 24hr post-CHI.** Group 1 cytokines (IL-1β, IL-13, and KC) and MAPK phospho-proteins (phospho-Atf2 and phospho-Mek1/2) (red) co-stained with GFAP (green) and DAPI (blue) showed minimal astrocyte co-localization in the hippocampus 24h post-CHI in female 3xTg mice aged 2-4mo (scale bar: 500µm).

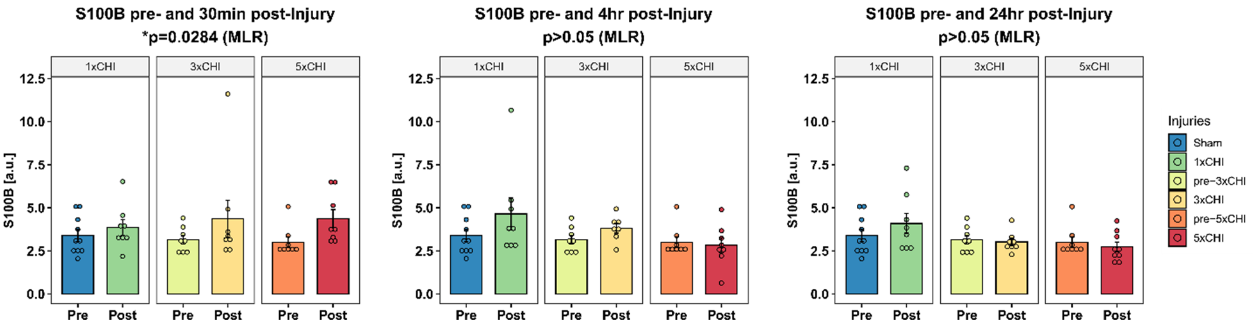


**Supplementary Figure S14. Cortical astrocytes become reactive following mild TBI: S100B.** Cortical S100 calcium-binding protein B (S100B) is significantly upregulated 30min post-injury vs pre-injury but not 4hr or 24hr post-injury (*p<0.05; multiple linear regression model, pre/post and number of injuries) (mean ± SEM).

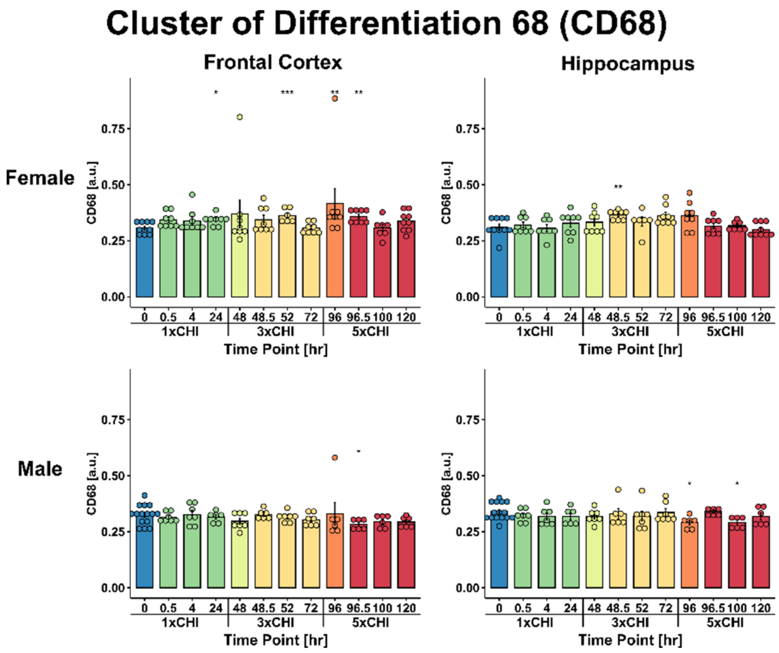


**Supplementary Figure S15.** CD68 was measured by ELISA in male and female frontal cortex and hippocampus tissue (*p<0.05, **p<0.01, ***p<0.001, Wilcox test vs sham) (mean ± SEM). Male and female samples run simultaneously.
